# The rhomboid-like pseudoprotease TMEM115 defines a two-factor mechanism for Rab6A effector recruitment

**DOI:** 10.64898/2026.09.07.749849

**Authors:** Boyan Zhang, Owain Bryant, Nadine Muschalik, Clémence Levet, Laura Charlier, Ángela Moncada-Pazos, Nina Jajčanin-Jozić, Xin Shen, Fangfang Lu, Abhimanyu Gowda, Stefan Düsterhöft, Susan Lea, Sean Munro, Matthew Freeman

**Affiliations:** Sir William of Dunn School, University of Oxford, Oxford, UK; Structural Biology, St Jude Children’s Research Hospital, Memphis, Tennessee USA; National Cancer Institute, National Institutes of Health, Frederick, Maryland, USA; Division of Cell Biology, MRC Laboratory of Molecular Biology, Cambridge, UK; Institute of Molecular Pharmacology, RWTH Aachen University, Aachen, Germany; Genbioma Aplicaciones, Navarra, Spain; HMU Health and Medical University, Campus Düsseldorf/Krefeld, Germany

## Abstract

Protein trafficking and organelle identity depends on precise localisation of membrane proteins, anchored by small GTPases that bind to specific cellular compartments. Here we show that the rhomboid-like pseudoprotease TMEM115 defines an unexpected two-factor system with Rab6A to target the golgin TMF1. In the absence of TMEM115, TMF1 is delocalised from the rim of the Golgi apparatus; instead, it is retained in Golgi-targeting vesicles. TMEM115, TMF1, and Rab6A assemble into a ternary complex which, in striking contrast to canonical rhomboid-like mechanisms, is independent of transmembrane domain interactions. Disruption of the TMEM115-Rab6A-TMF1 complex affects the efficiency of trafficking of proteins from the Golgi; moreover, human disease-associated mutations target the interfaces of this ternary complex, thereby delocalising TMF1 from the Golgi. Finally, loss of TMEM115 in *Drosophila* causes abnormal Golgi morphology, and mouse knockouts undergo perinatal death. We propose that two-factor effector recruitment defines a mechanism for precise Rab GTPase-mediated membrane targeting.

## Introduction

The eukaryotic cell comprises distinct membrane compartments, and the machinery of membrane trafficking has evolved to ensure that proteins are targeted precisely. The rhomboid-like superfamily of polytopic membrane proteins exemplifies this spatial control. Across the superfamily, members have well-defined and functionally important locations in specific membranes. For example, the rhomboid protease RHBDL4 is confined to the endoplasmic reticulum (ER), RHBDL2 is primarily at the plasma membrane, and RHBDL1 and RHBDL3 are located in the Golgi apparatus and endosomes, respectively (Bergbold and Lemberg, 2013). As well as the rhomboid proteases, the superfamily includes multiple rhomboid-like pseudoproteases that have lost their catalytic sites through evolution. Some of these, like iRhoms and derlins, are well characterised, but others remain obscure. We have explored the role of a poorly characterised, Golgi-localised rhomboid-like protein, TMEM115 (Ivanova et al., 2008; Ong et al., 2014), one of the most highly conserved members of the rhomboid superfamily.

The Golgi apparatus is the cellular hub for processing, sorting, modifying and recycling proteins and lipids. Nearly all proteins that are destined either for secretion or membrane localisation pass through the Golgi, arriving in vesicles from the ER, and then trafficked onwards to the later secretory pathway. The Golgi also contains resident proteins, needed to perform its multiple functions including, for example, glycosyltransferases; these must be precisely retained in their correct locations. This requirement to combine vesicle trafficking, maturation and sorting of a broad range of secreted and membrane proteins, while maintaining its own specialised proteome, means that spatial control of proteins in the Golgi is particularly complex.

One important class of resident Golgi proteins are the golgins. These well-conserved large coiled-coil proteins (at least 11 in mammals) are anchored to the cytoplasmic face of the Golgi and form a ‘tentacular’ mesh to capture cargo-containing vesicles (Witkos and Lowe, 2016). Individual golgins selectively capture incoming vesicles via their membrane-distal N-termini (Wong et al., 2017; Wong and Munro, 2014) and are themselves anchored to the Golgi membrane via their C-termini, usually through transmembrane domains (TMDs), or by binding to small membrane-associated GTPases of the Arl, Arf, and Rab families.

This function of Rab GTPases in locating golgins exemplifies the wider function of Rabs in targeting membrane-associated proteins throughout the cell. More than 60 Rabs have been identified in the human genome (Banworth and Li, 2018). In their GTP-bound forms, Rabs bind and thereby recruit effector proteins to specific membranes. This model predicts that effector proteins reside where active Rabs are localised, but accumulating evidence shows that Rabs do not always precisely locate their effector proteins. For example, the Rab5 effector EEA1 (Simonsen et al., 1998) is located in cytoplasmic endosomes, whereas its effectors APPL1/2 are enriched beneath the PM and barely co-localise with EEA1 (Miaczynska et al., 2004). Notably, even APPL1 and APPL2 only partially co-localise (Miaczynska et al., 2004). In another example, Rab6 is enriched in the Golgi apparatus and the *trans*-Golgi network (TGN), but its effectors (*e.g*. BICD1/2, TMF1, DCTN1) are sub-compartmentalized in distinct ways (Fridmann-Sirkis et al., 2004; Hoogenraad et al., 2001; Matanis et al., 2002; Short et al., 2002; Splinter et al., 2010). These examples make the point that while small GTPases like Rabs play an important role in locating their effector proteins, additional machinery must exist to ensure precise localisation of effectors.

In this study, we reveal a two-factor mechanism for precise targeting. We have discovered that the rhomboid-like pseudoprotease TMEM115 as an essential factor in the correct Golgi targeting of TMF1. TMEM115 forms a ternary complex with TMF1 and Rab6A, and without TMEM115, Rab6A alone cannot efficiently target TMF1. We extend this idea by showing that Rab6A is also inefficient at targeting its well-established effector BICD2, and that Rab5A is inefficient at targeting EEA1. Consistent with its role in locating TMF1, cells lacking TMEM115 are defective in protein secretion. At the organismal level, TMEM115 is essential for maintaining Golgi structure in flies, and its loss compromises postnatal mouse viability. We propose that the prevailing Rab-effector targeting model needs adjusting, highlighting that Rabs are not always sufficient, and that cofactors ensure the fidelity and efficiency of protein targeting.

## Results

### TMEM115 belongs to the rhomboid protein family and is enriched on the Golgi rim

Although sequence analysis suggests that TMEM115 is a distant member of the rhomboid superfamily (our analysis below and InterPro (Blum et al., 2021)), the results of a previous study challenged this notion by reporting that TMEM115 contains four TMDs (Ong et al., 2014). No *bona fide* rhomboids are known to have fewer than six TMDs (Bergbold and Lemberg, 2013; Dulloo et al., 2019; Freeman, 2014). To resolve this discrepancy, we examined the topology of TMEM115 by epitope-tagging the predicted loops between TMDs (**Figs. 1A, 1B, S1B**). By permeabilising cells with either 0.2% Triton X-100 or digitonin, which respectively permeabilises all cellular membranes or only the plasma membrane, we were able to determine by antibody staining whether each tagged loop resides in the lumen or the cytoplasm (**Fig. S1A**). As a control for the degree of permeabilisation, an antibody that targets the Golgi luminal domain of TGN46 could only detect TGN46 in cells permeabilised by Triton X-100 but not digitonin (**Fig. S1B**). The introduction of Flag or Myc epitopes into the five predicted loops and the N- and C-termini provided a clear result: their accessibility implied that, consistent with its predicted identity as a member of the rhomboid-like superfamily, TMEM115 contains six TMDs, with a short N-terminal and a long C-terminal cytoplasmic tail (**Figs. 1B, S1B**).

**Figure 1.**
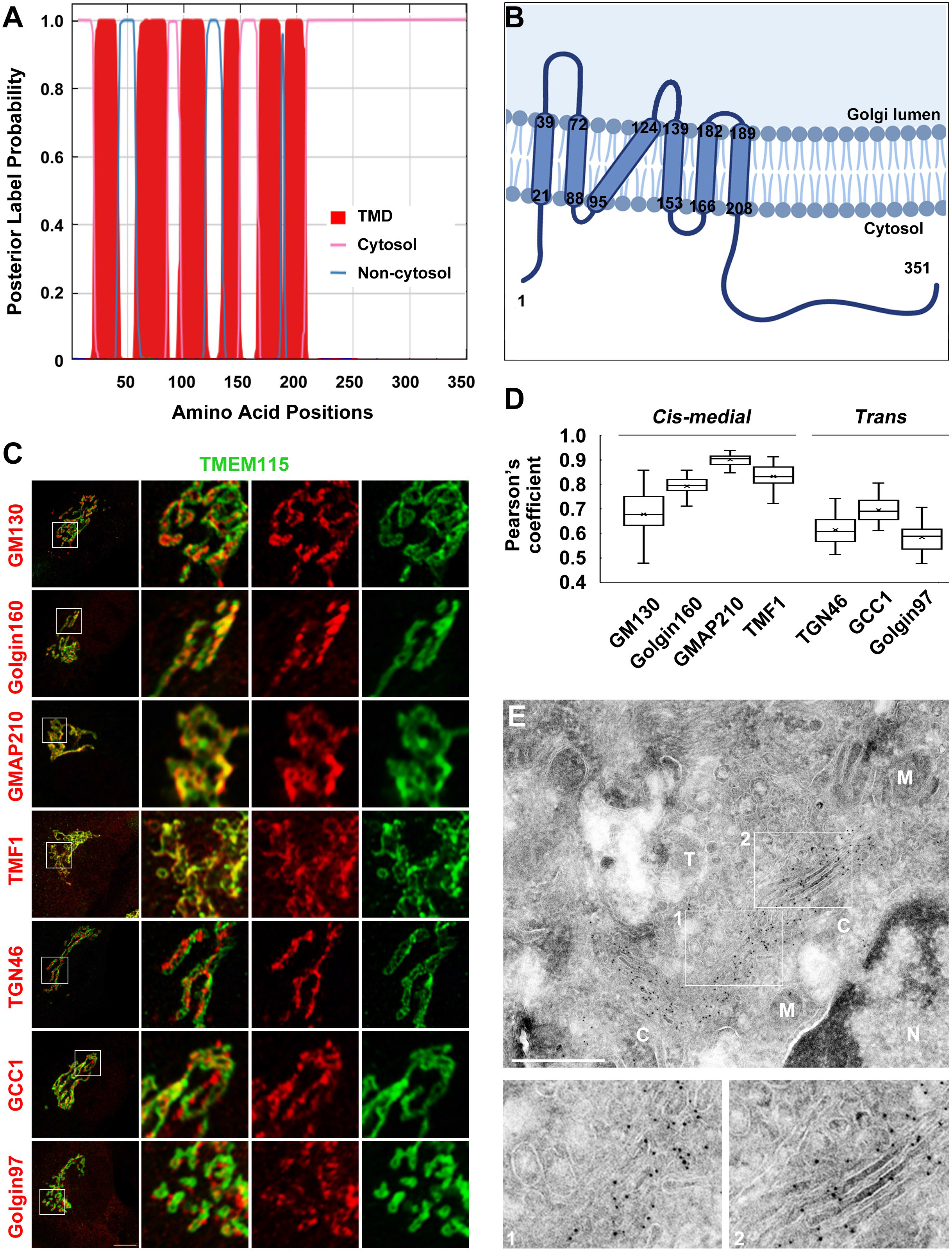
TMEM115 contains six TMDs and is enriched in the *cis-medial* Golgi. (A, B) Hydrophobicity analysis and the experimentally validated topology of TMEM115. The topology shown was analyzed by DeepTMHMM (A). The annotated TMD boundaries are derived from AlphaFold 3 (AF3) predictions and experimental validation (B). (C - E) TMEM115 is enriched on the Golgi rim in the *cis-medial* Golgi. (C) RPE1 cells stably expressing TMEM115-HA were imaged using Airyscan super-resolution microscopy. The boxed areas in the left panels are shown at higher magnification to the right. The Pearson’s coefficient of localisation of TMEM115 and the indicated golgin proteins are shown in (D). (E) RPE1 cells stably expressing TMEM115-HA were processed for immuno-transmission electron microscopy. Numbered boxes in the top panel are shown at higher magnification below. N, nucleus; M, mitochondria; C, cis-Golgi; T, trans-Golgi/TGN. Scale bar, 500nm. Throughout this study, unless otherwise indicated, microscopy experiments were done in RPE1 cells and scale bars indicate 10μm. Quantification of data was derived from at least 30 cells in each group, shown as box-and-whisker plot with mean and P values.

Using super-resolution Airyscan microscopy to localise TMEM115 in human RPE1 cells, we confirmed that it resides in the Golgi apparatus (Ivanova et al., 2008; Ong et al., 2014). In addition, we found that it is restricted to the rim of the Golgi (**Fig. 1C**), the location of active vesicle fusion and budding. This characteristic ring-like pattern in the Golgi was also observed in different cell types, such as COS7, Hela, MCF7, and MEFs (**Fig. S2A**). Co-localisation analysis showed that TMEM115 co-localises most closely with early-medial Golgi proteins such as GM130, GMAP210, Golgin160, and TMF1, but not the late Golgi proteins like GCC1, Golgin97, and TGN46 (**Fig. 1D**). Lastly, we used ultrastructural immuno-electron microscopy to locate TMEM115 and found that, consistent with the immunofluorescent results, C-terminally HA-tagged TMEM115 was primarily localised in the early-medial Golgi compartments as well as Golgi-associated vesicles, but not the TGN (**Fig. 1E**).

### TMEM115 is required for the Golgi localisation of TMF1

Using tandem mass tag quantitative mass spectrometry (TMT-MS), we compared the proteome of isolated Golgi membranes from control and TMEM115 siRNA knockdown (KD) RPE1 cells by. Combining the results from two different siRNAs, a total of 326 proteins were significantly altered (*P* < 0.05). Among these hits, we measured a change in abundance of greater than 50% in 4/178 of down-regulated and 26/148 of up-regulated proteins (**Figs. 2A, S2B**). Other than TMEM115 itself, the protein showing the greatest decrease was the golgin TMF1. We validated this result in TMEM115 knockout cells (2 independent lines named clone 1 and 6, hereafter called KO1 and KO6) and KD cells (**Figs. 2B, 2C, S3A – S3E**).

**Figure 2.**
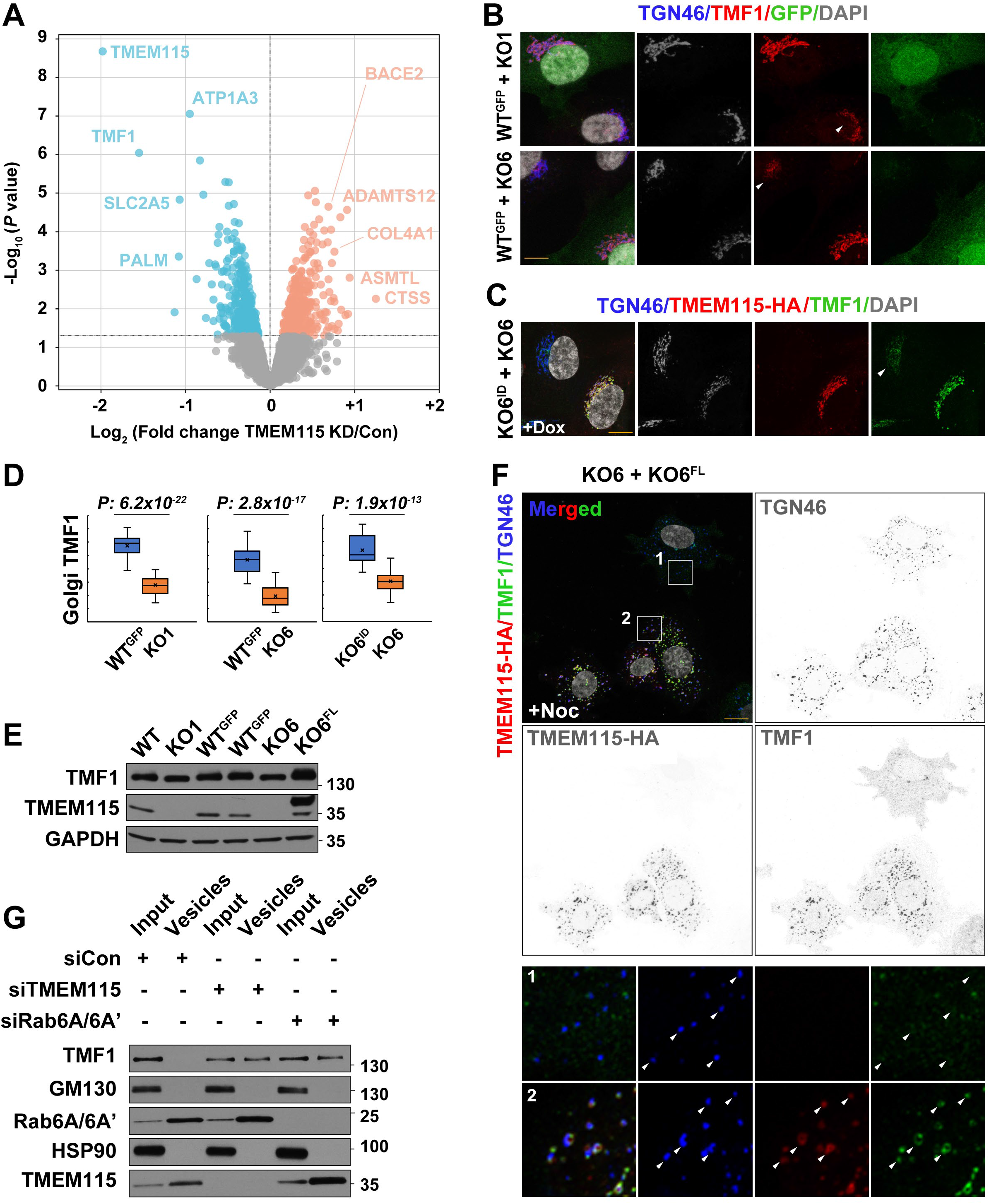
TMEM115 is required for the Golgi localisation of TMF1. (A) Effect of TMEM115 knockdown on the Golgi proteome. Golgi membranes were isolated from the TMEM115 knockdown and control cells, followed by TMT-mass spectrometry. The plot is derived from two TMEM115 knockdown datasets, each performed in triplicate. (B - E) De-localisation of TMF1 by knockout (KO) of TMEM115 can be rescued by TMEM115 expression. In (B) WT^GFP^ and TMEM115 KO1 or KO6 cells were co-cultured; in (C) KO6^ID^ (in which inducible TMEM115-HA was stably expressed in the KO6 cells) and KO6 cells were co-cultured for 10 hours in the presence of doxycycline. The intensity of fluorescent staining of Golgi TMF1 is shown in (D). (E) The total protein level of TMF1 was not affected by TMEM115 KO. Arrowheads in (B) and (C) indicate TMEM115-negative cells in the co-cultures. (F) Apparently Golgi-retained TMF1 in TMEM115 KO cells is not actually integrated in the Golgi. KO6 and KO6^FL^ cells were co-cultured and treated with nocodazole for 3h. In TMEM115 KO cells, nocodazole treatment (+Noc) resulted in a decrease of TMF1 in Golgi mini stacks and an increase of cytoplasmic TMF1. Numbered boxes are shown at higher magnification in the lower panels. Arrowheads, Golgi mini stacks. (G) In the absence of TMEM115 or Rab6A,TMF1 is enriched in vesicles compared to controls. Total vesicles were isolated from cells knocked down for TMEM115 or Rab6A.

Fluorescent microscopy showed that the loss of TMEM115 resulted in Golgi fragmentation and reduced Golgi-localised TMF1 (**Figs. 2B – 2D, S3D, S3D, S3E**), and that both phenotypes could be rescued by expressing full-length TMEM115 (**Figs. 2C, 2D**). Western blotting for TMF1 showed that despite the reduced staining in the Golgi, the total level of TMF1 in the cell was unaffected by the loss of TMEM115 (**Fig. 2E**), suggesting that TMF1 is de-localised from the Golgi rather than being degraded.

Both TMEM115 and TMF1 are widely conserved across eukaryotic evolution. We therefore examined whether their functional relationship was also conserved. Deleting the *Drosophila* homologue of TMEM115, *CG9536*, in flies expressing endogenously GFP-tagged Tmf resulted in a decrease of the GFP signal in the Golgi of larval salivary glands, but no decrease of the total protein level of GFP-Tmf (**Figs. S2C – S2E**). This phenotype in whole organisms mirrors our observations in mammalian cells.

### TMEM115 and Rab6A together target TMF1 to Golgi membranes

We noticed in mammalian cells that, although loss of TMEM115 substantially decreased TMF1 at the Golgi, a residual level always remained. TMF1 does not contain a TMD but is instead membrane-anchored by Rab6A, deletion of which showed a similar partial loss of TMF1 Golgi localisation (Yamane et al., 2007). To investigate whether TMEM115 and Rab6A are responsible for distinct Golgi pools of TMF1, we compared TMF1 localisation in TMEM115 and Rab6A/6A’ knockdown cells individually or in combination (Rab6A has two different isoforms and in all experiments we knocked down both). Individual knockdown of each de-localised Golgi TMF1 to a similar extent, and double knockdown did not further decrease residual Golgi TMF1 (**Figs. S3A – S3E**). We therefore conclude that TMEM115 and Rab6A regulate the same pool of TMF1 in the Golgi rather than being responsible for distinct sub-populations.

A possible cause for the residual TMF1 staining around the Golgi is that, instead of being Golgi-anchored, it might represent a fraction associated with Golgi-proximal vesicles that cannot be resolved by confocal microscopy. To test this, we disrupted vesicle trafficking by depolymerising cellular microtubules with nocodazole. This led to almost complete loss of detectable TMF1 associated with Golgi ministacks in the KO6 cells. In KO6 cells rescued with full-length TMEM115 (KO6^FL^), the Golgi ministack localisation of TMF1 was maintained (**Fig. 2F**). Meanwhile, TMF1 staining associated with the cytoplasm was increased in KO6 cells, but not in the rescued KO6^FL^ cells (**Fig. 2F**). These data suggest that the residual TMF1 labelling on the Golgi seen upon loss of either TMEM115 or Rab6A is not in fact Golgi-associated; instead, it is in Golgi-proximal vesicles that are dispersed when microtubules are depolymerised. This interpretation is further confirmed biochemically: when either TMEM115 or Rab6A was knocked down, TMF1 became enriched in vesicles, whereas in control cells, TMF1 was undetectable in the vesicle fraction (**Fig. 2G**).

Overall, these results demonstrate that both TMEM115 and Rab6A are required for Golgi membrane association of TMF1. In the absence of either, TMF1 dissociates from the Golgi apparatus and becomes accumulated in neighbouring vesicles.

### TMEM115 and TMF1 interact through their C-termini

Previous reports point to Rab6A alone being responsible for Golgi localisation of TMF1 (Fridmann-Sirkis et al., 2004; Yamane et al., 2007). We therefore first investigated whether TMEM115 regulates the level, localisation, or activity of Rab6A. In TMEM115 KO Golgi extracts, the protein levels of Rab6A were unaffected (**Fig. S3F**). TMEM115 KD did not affect the Golgi localisation of GFP-Rab6A (**Figs. S3G, S3H**), nor did overexpression of TMEM115 in KO6 cells increase Golgi staining of Rab6^GTP^ (**Figs. S3I, S3J**). Consistent with this, our Golgi proteomics data showed little difference between KD and control cells in the levels of other known Rab6A effectors (GCC2, BICD1/2, MYH9/10, OCRL, ERC1, DYNLRB1) (G. Dornan and C. Simpson, 2023), or the Rab6A regulators (RIC1, Rab33B, RGP1, and GDI1/2), or Rab6A itself. We conclude that TMEM115 does not regulate the level, localisation, or activity of Rab6A.

We next asked whether physical interaction between TMEM115 and TMF1 underpins their functional relationship. Immunoprecipitation (IP) showed that TMEM115-HA binds to TMF1 and Rab6A but not to several other soluble golgins or Golgi-associated Rabs (**Fig. 3A**). Reciprocally, GFP-TMF1 pulled down endogenous TMEM115 and Rab6A (**Fig. 3B**). To define the domains mediating the TMEM115, TMF1, Rab6A interactions, we stably expressed a series of TMEM115 truncations in the KO6 cells. We found that whereas residues 1-270 of TMEM115 are required for its Golgi localisation, only the predicted cytoplasmic helix of TMEM115, comprising residues 281-300, is needed for its interaction with TMF1 and Rab6A (**Figs. 3C, 3D, S4**). Expressing constructs containing this domain could also restore the Golgi localisation and eliminate the vesicle accumulation of TMF1 in KO6 cells (**Fig. S5**).

**Figure 3.**
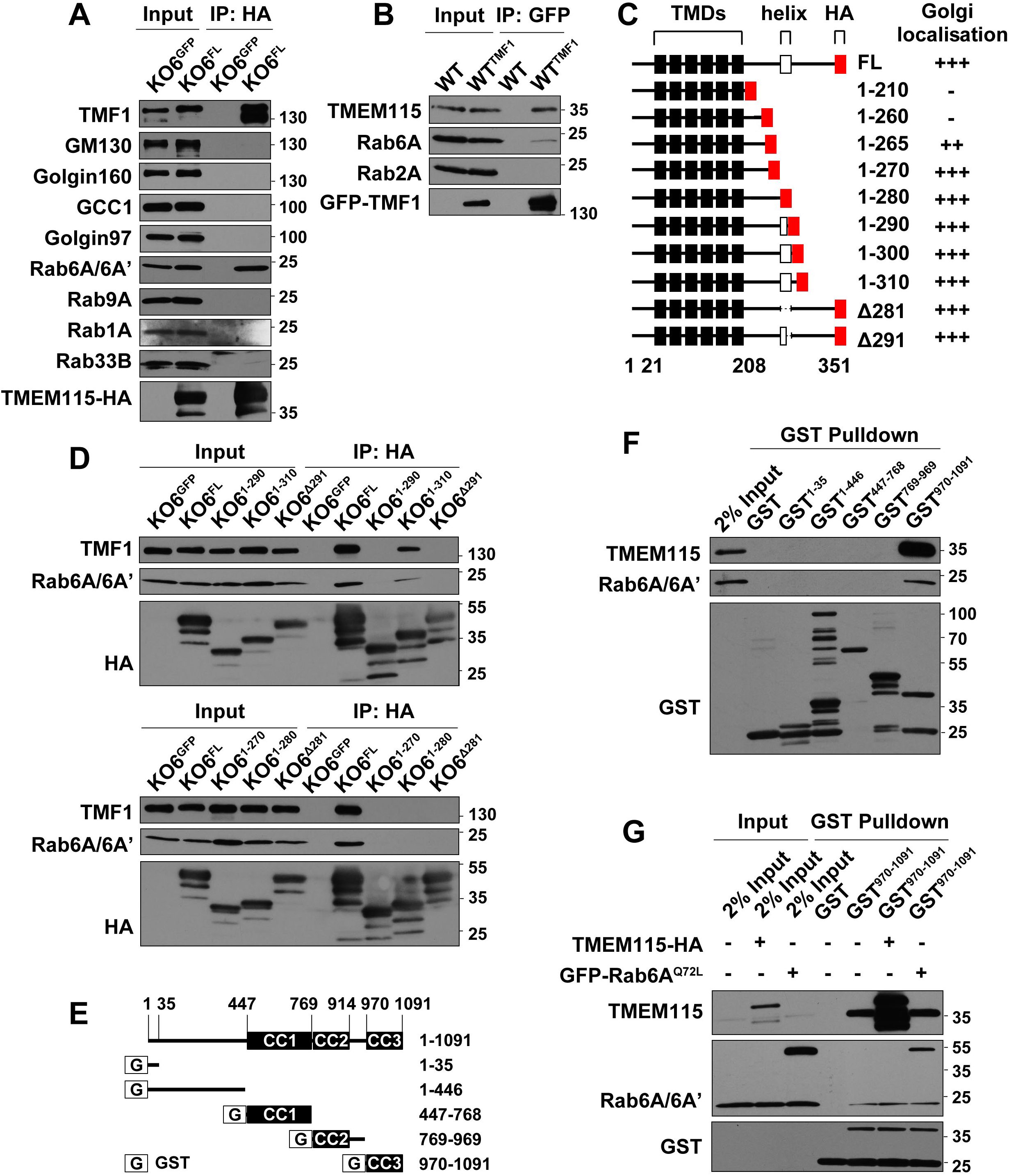
The cytoplasmic helix of TMEM115 and the Golgi proximal C-terminal of TMF1 mediate the TMEM115-TMF1-Rab6A interaction. (A, B) TMEM115 interacts with TMF1 and Rab6A. (A) Immunoprecipitation from KO6 and KO6^FL^ cells. (B) Immunoprecipitation from WT and stable GFP-TMF1- expressing cells. (C, D) The cytoplasmic helix of TMEM115 mediates its interaction with TMF1 and Rab6A. (C) TMEM115 deletion constructs used in the experiment. The Golgi localisation of each construct is summarized to the right. (D) Immunoprecipitation from KO6 cells stably expressing the indicated TMEM115 constructs. (E, F) The C-terminus of TMF1 interacts with TMEM115 and Rab6A. (E) TMF1 constructs used in the experiment; CC, coiled coil. (F) GST or GST-tagged TMF1 fragments were used as baits, pulling down proteins from total cell lysates. (G) TMEM115 and Rab6A do not compete for TMF1. GST or GST-TMF1^970-1091^ proteins were used as baits, pulling down proteins from cells transiently over- expressing TMEM115-HA or GFP-Rab6A^Q72L^.

**Figure 4.**
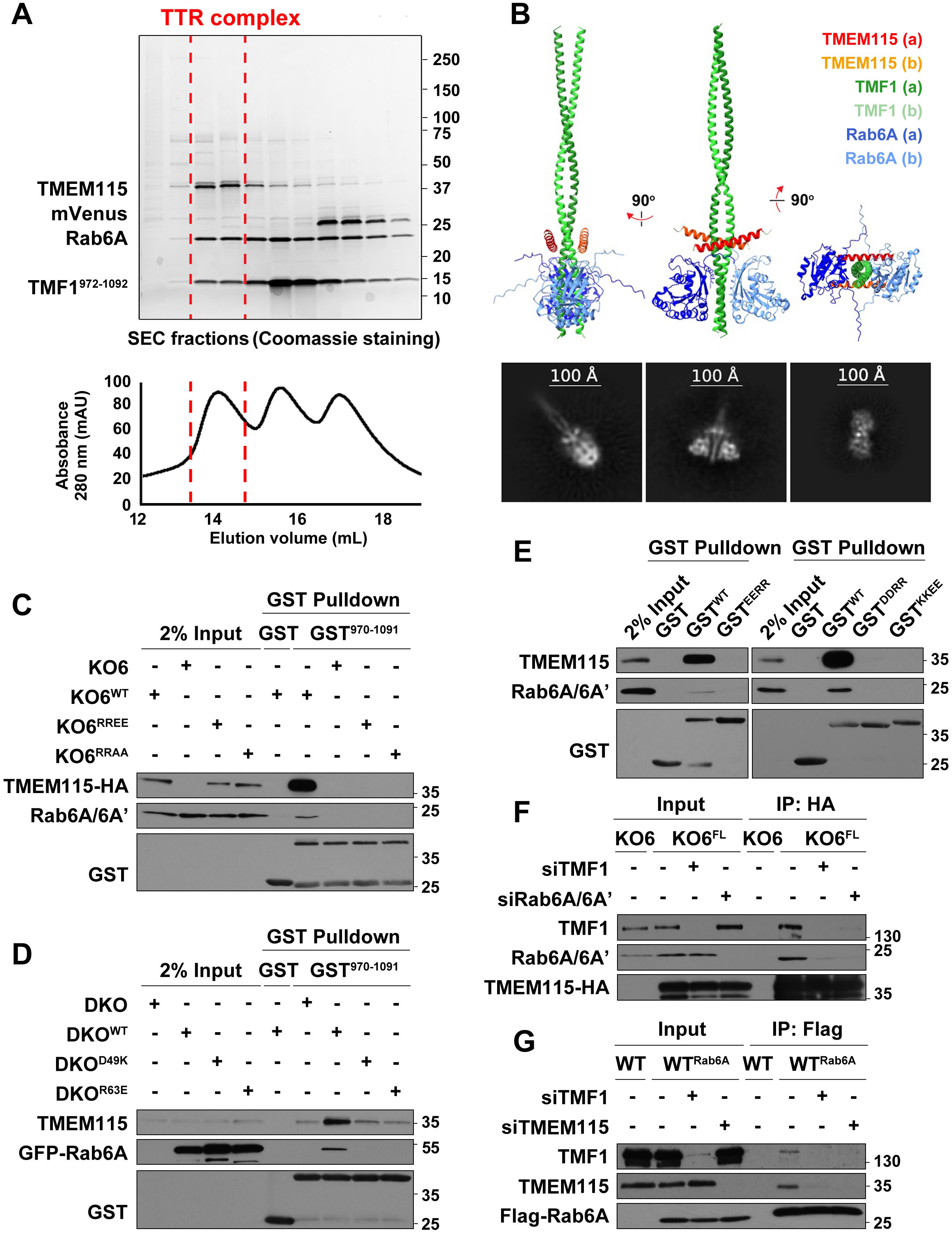
TMEM115, TMF1, and Rab6A form a complex. (A) TMEM115, TMF1, and Rab6A form a complex *in vitro*. The eluant from a mixture of human full-length TMEM115, Rab6A, and TMF1 (residues 972-1093) in LMNG detergent was fractionated on an S6 10/300 size-exclusion column (SEC). Proteins present in fractions were analyzed by SDS-PAGE and Coomassie staining; mVenus was derived from cleavage of the TMEM115-mVenus-twin-Strep fusion protein. (B) AlphaFold 3-predicted structure of the TMEM115 (residues 275-305)-TMF1 (residues 970-1091)-Rab6A ternary complex. Representative 2D class averages from Cryo-EM data are shown below each view of the model. (C, D) TMEM115 and Rab6A are both essential for forming the TTR complex. The indicated GST and WT GST-TMF1^970-1091^ proteins were used as bait, pulling down proteins from total cell lysates of the TMEM115 KO, TMEM115-HA WT or mutant- expressing (KO6, KO6^WT^, KO6^RREE^ or KO6^RRAA^) cells, or the Rab6A/6B DKO, GFP- Rab6A WT or mutant-expressing (DKO, DKO^WT^, DKO^D49K^ or DKO^R63E^) cells. (E) The TMEM115-TMF1 and TMF1-Rab6A interfaces are both essential for formation of the TTR complex. GST coupled to WT or mutant GST-TMF1^970-1091^ proteins were used as bait, pulling down proteins from total cell lysates. (F, G) TMEM115, Rab6A, and TMF1 are mutually required for the TTR complex formation *in vivo*. Cells stably expressing Flag-Rab6A (WT^Rab6A^) or TMEM115-HA (KO6^FL^) were knocked down for the indicated genes, followed by an IP assay.

In a complementary set of experiments, we mapped the part of TMF1 associates with TMEM115. We found that all the three coiled-coils spanning residues 447-1091 of mouse TMF1 are simultaneously required for its Golgi localisation (**Fig. S6**). We therefore used *in vitro* GST-pulldown assays, allowing us to test even those constructs that were not Golgi-localised *in vivo*. The GST-tagged C-terminus of mouse TMF1 (residues 970-1091, hereafter referred as GST-TMF1) was the only fragment that pulled down endogenous TMEM115 and Rab6A (**Figs. 3E, 3F**). Notably, increasing the amount of TMEM115 or Rab6A by over- expressing TMEM115-HA or GFP-Rab6A^Q72L^ did not reduce the interaction of TMF1 with the other partner (**Fig. 3G**), indicating that TMEM115 and Rab6A do not compete for binding to TMF1. Together, these results demonstrate that the short cytoplasmic helix (residues 281- 300) of TMEM115 and the Golgi proximal end (residues 970-1091) of TMF1 mediate the interaction between them, and with Rab6A

### TMEM115, TMF1, and Rab6A form a ternary complex *in vitro*

To investigate the potential of TMEM115, TMF1, and Rab6A to form a complex, we mixed *in vitro* recombinant human proteins: full-length Rab6A loaded with non-hydrolysable GTP (GTPγS), full-length TMEM115, and the TMEM115-interacting fragment of TMF1 (human residues 972-1093, corresponding to mouse residues 970-1091). Fractionation by size-exclusion chromatography (SEC) of the reconstituted complex produced higher-order species with a distinct elution profile containing all three proteins (**Fig. 4A**). We assessed the suitability of this ternary complex (which we refer to as the TTR complex) for structural determination by cryo-EM but found that the mobility of the TMEM115 transmembrane module with respect to the cytoplasmic domain prevented structural solution (**Figs. S7A**).

To overcome this constraint, we reconstituted a ternary complex in which full-length TMEM115 was replaced by a fragment comprising its residues 275-305, which includes the binding helix (residues 281-300). In cryo-EM, this version of the complex did provide two-dimensional class averages that showed views of an elongated structure (‘side views’) and averages of an alternative view (‘top views’, **Figs. 4B, S7B, S7C**). The number of different views was limited by preferred orientations in vitreous ice and, combined with the small size of the complex, this prevented successful three-dimensional reconstructions. Nevertheless, the two-dimensional class averages were sufficient to demonstrate consistency with a highly confident AlphaFold 3 prediction of the complex (**Figs. 4B, S7D**), wherein two copies of human TMF1 (residues 972-1093) form a parallel alpha-helical right-handed coiled-coil, and two copies of the TMEM115 cytoplasmic helix (residues 275-305) sandwich the TMF1 coiled-coils, adjacent to the Rab6A binding sites. The inferred structure also shows that both copies of TMF1 participate in Rab6A binding, with two Rab6A molecules binding at opposite sides of the TMF1 dimer.

Since the Rab6A-TMF1 binding mode resembles that of Rab6A in complex with the golgin GCC2 (PDB: 3BBP, (Burguete et al., 2008)) or the kinesin protein Kif20A (PDB: 5LEF, (Miserey-Lenkei et al., 2017)) (**Fig. S8A**), we tested whether Rab6A and TMF1 (residues 972-1093) could form a complex on their own. SEC showed that Rab6A and TMF1 (residues 972-1093) formed a complex in the absence of TMEM115. In contrast, neither Rab6A and TMEM115 (residues 275-305), nor TMF1 (residues 972-1093) and TMEM115 (residues 275- 305) interacted in the absence of the third protein (**Figs. S8B, S8C**). These results indicate that the affinity between Rab6A and TMF1 can drive their interaction when they are at sufficiently high concentrations *in vitro*.

### *In vivo*, TMEM115 is essential for formation of the TTR complex

As is common with membrane proteins, AlphaFold 3 predictions of full-length TMEM115 in complex with TMF1 and Rab6A positioned the TMDs in configurations incompatible with a membrane-embedded state. Importantly, though, this mispositioning of the TMDs had no effect on the predicted TMEM115-TMF1 and TMF1-Rab6A interfaces. At the TMEM115- TMF1 interface, Arg282 and Arg283 of human TMEM115(a) are predicted to interact with Glu1061 and Glu1064 of human TMF1(a) (equivalent to Glu1059 and Glu1062 of mouse TMF1), while Arg294 of TMEM115(a) contacts Glu1061 of human TMF1(b) (equivalent to Glu1059 of mouse TMF1), with additional hydrophobic contributions (**Fig. S9A**). At the TMF1-Rab6A interface, Glu1068, Asp1072, and Asp1075 of human TMF1(a) (equivalent to Glu1066, Asp1070, and Asp1073 of mouse TMF1) interact with Arg63 of Rab6A(a), and Lys1077 and Lys1081 of human TMF1(b) (equivalent to Lys1075 Lys1079 of mouse TMF1) interact with Asp49 of Rab6A(b), also supported by hydrophobic interactions (**Fig. S9B**).

These interfaces are arranged symmetrically around the TMF1 coiled-coil, with TMF1 acting as a bridge between TMEM115 and Rab6A. Notably, the residues predicted to mediate these interactions are conserved across eukaryotes (**Fig. S9C**), underscoring their likely functional significance.

We used site-directed mutagenesis to investigate the contribution of the predicted interfaces to the TTR complex. The WT or Arg282/Arg283 double mutant (where the Arginine was mutated to oppositely charged or neutral residues) TMEM115 was stably expressed in KO6 cells (KO6^WT^, KO6^RREE^ or KO6^RRAA^). GST-TMF1 pulldown assay showed that the TMEM115 mutations abolished not only the TMEM115-TMF1 interaction, but also the assembly of the TTR complex (**Fig. 4C**). We next tested a Rab6A/6B double mutant background (DKO)(Homma et al., 2019), which removes the possibility of compensation of the Rab6A KO by Rab6B, a Rab6 variant that has been proposed to be a TMF1-dedicated Rab GTPase (Fridmann-Sirkis et al., 2004). In these cells, the TTR complex was only assembled when WT GFP-Rab6A (DKO^WT^), but not GFP-Rab6A mutants (D49K and R63E for TMF1 interactions; DKO^D49K^ or DKO^R63E^) was present (**Fig. 4D**). Probing the other side of the predicted interactions, mutating mouse TMF1 residues Glu1059 and Glu1062 (GST^EERR^), Asp1070 and Asp1073 (GST^DDRR^), or Lys1075 and Lys1079 (GST^KKEE^) into oppositely charged residues all abolished the TTR complex (**Fig. 4E**). Finally, knocking down any component of the TTR complex disrupted the interaction of the others (**Figs. 4F, 4G**). Together, these results confirm the predicted interfaces and demonstrate that, *in vivo*, TMF1 and Rab6A do not interact in the absence of TMEM115.

### Human disease-associated mutations disrupt the TTR complex and delocalise TMF1 from the Golgi

Residues Asp1072 and Asp1075 of human TMF1 (equivalent to mouse TMF1 Asp1070 and Asp1073) are mutated in human large intestine and endometrioid carcinoma tissues, respectively (**Fig. 5A**). A nonsense mutation of the Arg63 residue of Rab6A is detected in the human pancreas carcinoma tissue (**Fig. 5A**). Furthermore, human variants were found at the Glu1061, Glu1068, and Asp1075 residues of TMF1 (equivalent to mouse TMF1 Glu1059, Glu1066, and Glu1073), and the Arg282 and Arg294 residues of TMEM115 (**Fig. 5A**). All these residues are critical for mediating the TMF1-TMEM115 and TMF1-Rab6A interactions, and those genomic variants are predicted to be deleterious or pathogenic (Cheng et al., 2023; Rentzsch et al., 2019; Rives et al., 2021). We found that the human disease-associated mutants, genomic variants, and our experimental mutations all disrupted TTR complex formation *in vivo*. In addition, except for the Asp1075Tyr mutation, all delocalised TMF1 from the Golgi (**Figs. 5A – 5C, S10A – S10D**).

**Figure 5.**
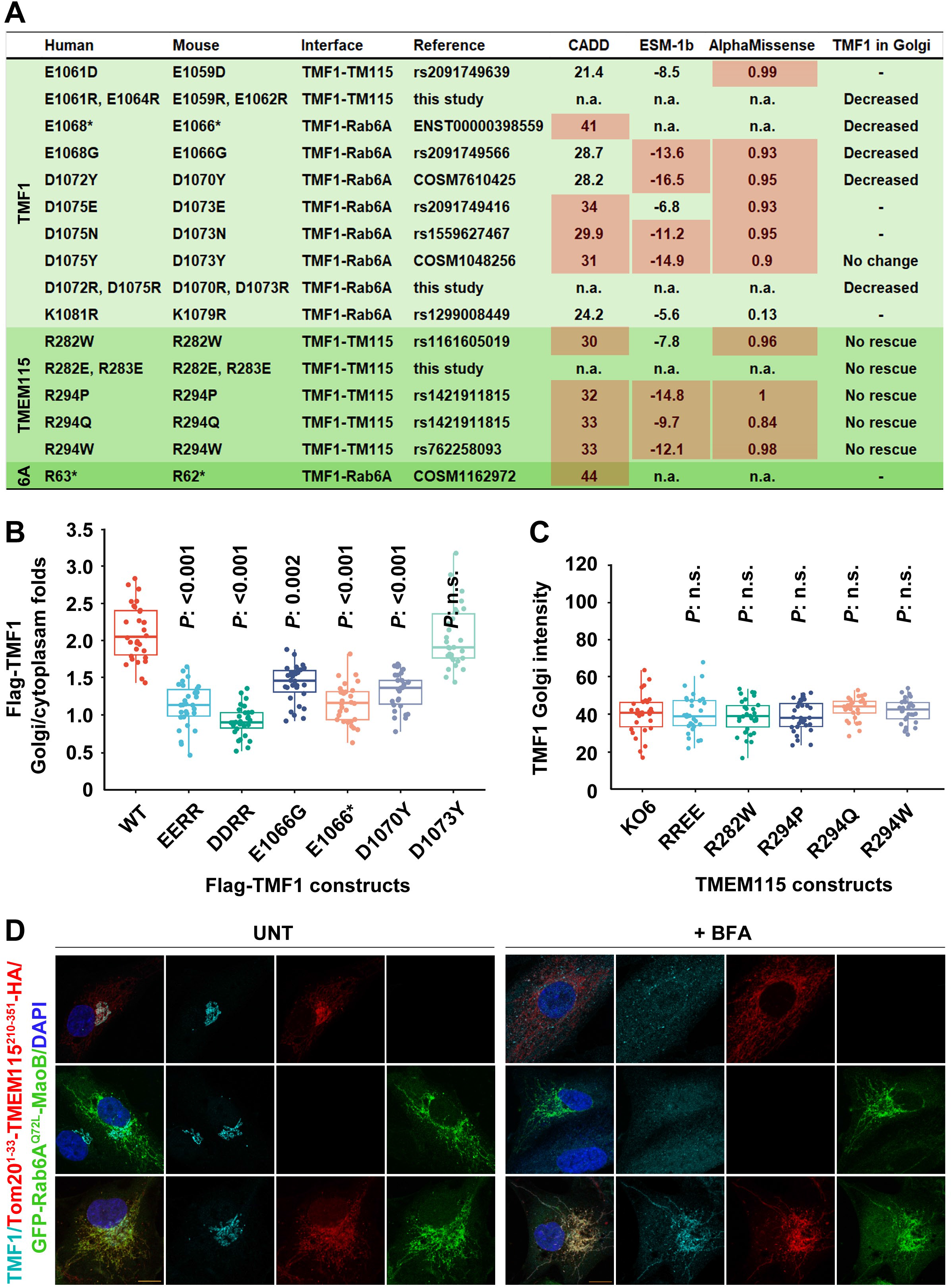
Formation of the TTR complex is required for the Golgi localisation of TMF1. (A - C) Mutations that disrupt the TMF1-TMEM115 or TMF1-Rab6A interfaces perturb TMF1 Golgi localisation. The human disease-associated mutations or genomic variants are derived from the Ensembl, NCI-TCGA, TOPMed, and COSMIC databases. (B) Cells were transiently transfected with Flag-tagged WT or mutant TMF1 constructs, and the intensity of Flag-TMF1 in the Golgi was normalized to that of the background staining of cytoplasm. (C) The KO6 and KO6 cells stably expressing TMEM115-HA mutants were co-cultured and Golgi TMF1 was quantified. Representative images for quantification analysis in (B, C) are in Figs. S10A, S10C. (D) TMEM115 and Rab6A synergistically locate TMF1. Cells stably expressing either Tom20^1-33^-TMEM115^210-351^-HA or GFP-Rab6A^Q72L^-MaoB alone, or both constructs together, were untreated (left panels) or treated with BFA for 1 hour (right panels).

### TMEM115 and Rab6A synergistically localise TMF1

To understand why both TMEM115 and Rab6A are needed to localise TMF1, we used the ‘mitochondrial anchor-away’ technique, in which a protein of interest is ectopically retargeted to the mitochondrial membrane, allowing one to assess whether it is sufficient to relocate its effectors (Wong and Munro, 2014). We replaced the last three residues required for prenylation of the GTP-locked Rab6A mutant (Rab6A^Q72L^) with the mitochondrial localisation signal of MaoB (Mitoma and Ito, 1992) (GFP-Rab6A^Q72L^-MaoB^492-520^). The cytoplasmic tail of TMEM115 (residues 210-315) was fused to the mitochondrial localisation signal of Tomm20 (Kanaji et al., 2000) (Tomm20^1-33^-TMEM115^210-351^-HA) (**Fig. S10E**). Upon stable expression of these relocalised proteins, neither alone, nor the two together, were able to relocate TMF1 to mitochondria (**Fig. 5D, left panels**).

We reasoned that this failure to retarget TMF1 might be caused by the stability of its Golgi localisation prior to expression of the ectopic proteins. To address this, we treated the cells with Brefeldin A (BFA), which causes Golgi collapse and thereby frees TMF1. In the presence of BFA, co-expressing both Tom20^1-33^-TMEM115^210-351^-HA and GFP-Rab6A^Q72L^- MaoB^492-520^ led to clear mitochondrial relocation of TMF1 (**Fig. 5D, right panels**). Significantly, even when expressed at a high level, GFP-Rab6A^Q72L^-MaoB^492-520^ alone was unable to mitochondrially target TMF1, or BICD2, another well-known Rab6A effector (**Figs. S11A, S11B**). These results imply that the inefficiency of Rab6A alone in localising effectors is not caused by expression levels; instead, TMF1 requires a combination of Rab6A and TMEM115 for efficient localisation.

### Rab5A is also inefficient at targeting its effector EEA1

The relatively low affinity of Rab6A towards effectors might cause its inefficiency (Noell et al., 2018). To probe whether this is confined to Rab6A, we also tested the efficiency of Rab5A, an endosomal Rab GTPase with a higher affinity for effectors (Mishra et al., 2010). Transiently expressed Rab5A^Q79L^-HA-MaoB^492-520^ was enriched in the mitochondria but was insufficient to relocate its well-defined effector EEA1 (Simonsen et al., 1998). Even in the presence of nocodazole, which releases EEA1-containing endosomes from microtubules, mitochondrial GTP-locked Rab5A only modestly recruited EEA1 (**Fig. S11C**). This second example suggests that the need for cofactors in the Rab-addressing system may be widespread.

### The TMEM115-TMF1 axis contributes to post-Golgi anterograde protein trafficking

The role of Rab6A in the secretory pathway is well established. To reveal the functional relationship between TMEM115 and TMF1 in the Golgi, we used TMT-MS to examine the Golgi proteome of RPE1 cells knocked down for TMF1. Across the results from using two different siRNAs, a total of 216 proteins were significantly altered (*P* < 0.05). The abundance of 2/104 of down-regulated proteins and 29/112 of up-regulated proteins were changed more than 50% (**Figs. 6A**, **S12A**).

**Figure 6.**
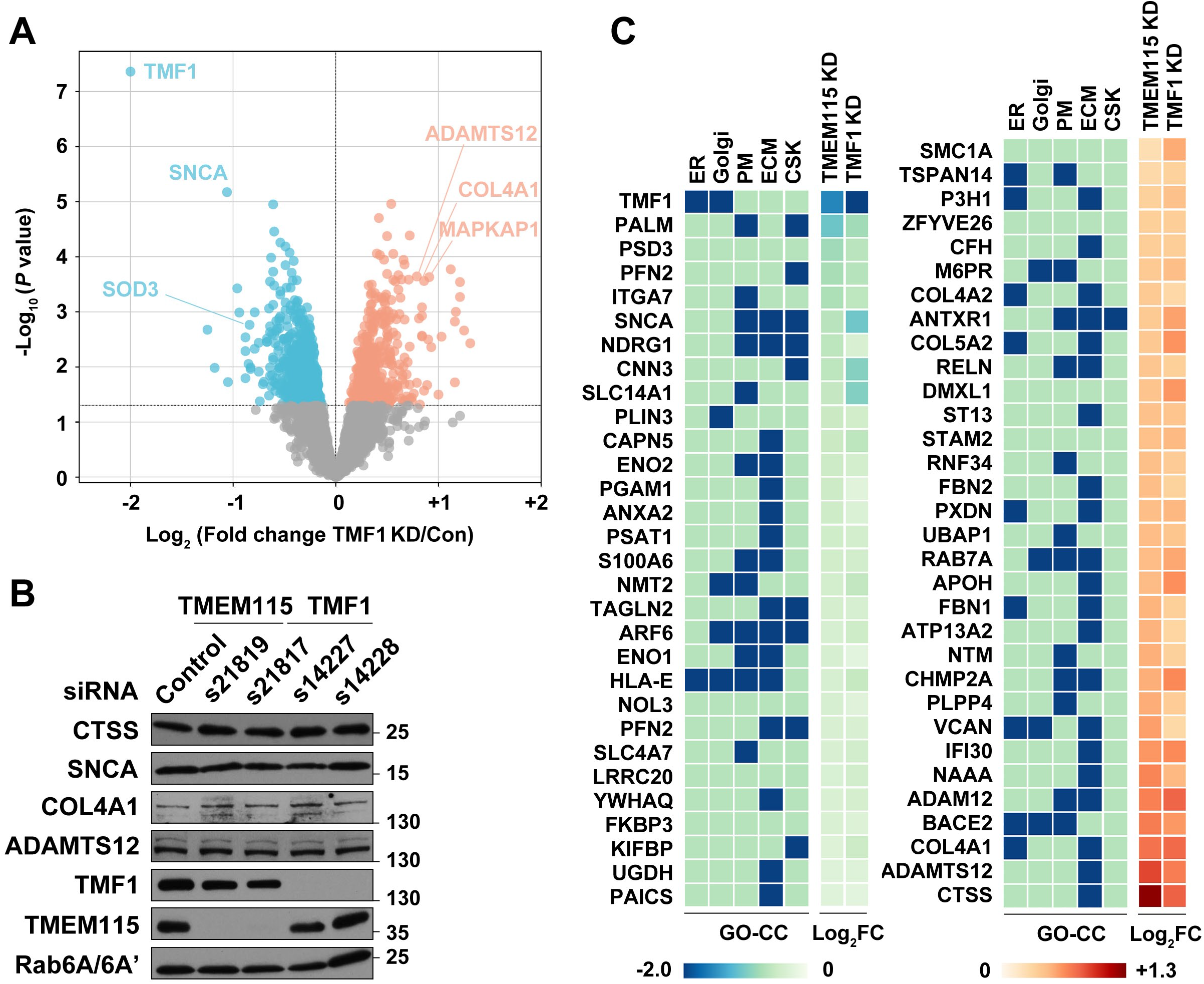
The TMEM115-TMF1 axis regulates post-Golgi protein trafficking. (A) Effect of TMF1 knockdown on the Golgi proteome. Golgi membranes were isolated from TMF1 knockdown and control cells, followed by TMT-mass spectrometry. The plot is derived from two TMF1 knockdown datasets, each in triplicate. (B) TMF1 and TMEM115 knockdowns do not affect total protein levels of a range of Golgi proteins. (C) The TMEM115-TMF1 axis affects the trafficking of secreted proteins. The common protein hits from TMEM115 and TMF1 knockdown Golgi proteomes across all analyses are grouped as up-regulated (left panel) and down-regulated (right panel). The GO subcellular distribution and the Log2(fold change of abundance) of these proteins are indicated by colour. CSK, cytoskeleton; ECM, extracellular matrix; ER, endoplasmic reticulum; PM, plasma membrane.

Comparing our results of knocking down TMEM115 (see above) and TMF1, we found that 29 proteins in common were upregulated, and 32 downregulated **(Figs. 7C, S12B)**. In the cases we examined, although clear changes were observed in Golgi abundance, there were only minor, if any, corresponding changes in overall protein levels (**Fig. 6B**). Gene Ontology Cellular Components (GO-CC) analysis revealed that most of the altered proteins are normally targeted to the plasma membrane or the extracellular matrix (ECM) (**Fig. 6C)**. Together, these results suggest the TMEM115-TMF1 axis functions in regulating post-Golgi anterograde trafficking, including secreted proteins destined for the ECM. This conclusion is consistent with a recently reported role of TMEM115 in cell migration (Sun et al., 2025; Yin et al., 2025).

**Figure 7.**
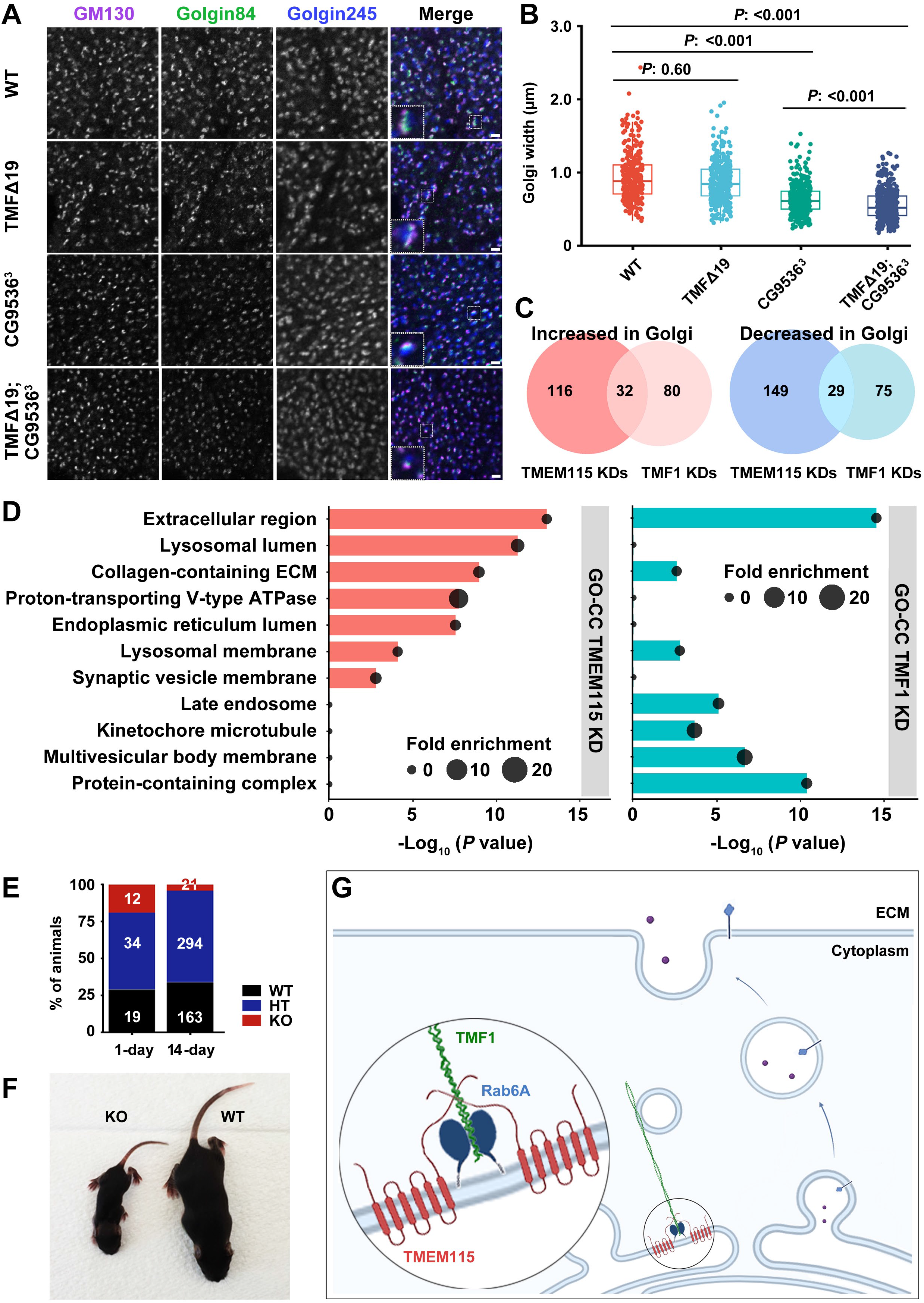
TMEM115 has a wider biological role than TMF1. (A, B) Deletion of the Tmem115 gene in *Drosophila*, *CG9536,* affects Golgi structure more than *Tmf* mutants. (A) Salivary glands of fly larvae were fixed and stained for the indicated cis (GM130)-to-trans (Golgin245) Golgi proteins. A single Golgi stack is shown magnified in right hand panels. Scale bar, 2µm. (B) The width of Golgi stacks was quantified. (C) TMEM115 knockdown affects Golgi proteome more than TMF1 knockdown. The consistently altered protein hits across two TMEM115 knockdown datasets and two TMF1 knockdown datasets are shown in a Venn diagram. More proteins are affected by TMEM115 knockdown than TMF1 knockdown. (D) TMEM115 and TMF1 knockdowns have common protein hits but also have distinct consequences. DAVID GO-CC analysis was applied to the consistently altered protein hits across two TMEM115 knockdown datasets and two TMF1 knockdown datasets. (E, F) TMEM115 KO mice primarily die postnatally. (E) Frequencies of the different genotypes at day 1 (n=65) and 14 (n=478) of life. (F) A representing image of the WT and KO littermates at 12 days of age. (G) Working model of the two-factor mechanism that localises TMF1 by the combined actions of TMEM115 and Rab6A in the Golgi apparatus, thereby enabling correct post-Golgi anterograde protein trafficking.

### TMEM115 has a broader biological function spectrum than TMF1

We finally examined the role of TMEM15 and TMF1 in whole organisms. In *Drosophila*, normal Golgi structure is indicated by the size and morphology of individual Golgi stacks. Knocking out *Tmf*, *Tmem115* (*CG9536)*, or both, did not change the polarity of Golgi stacks, but resulted in a significant decrease of their width **(Figs. 7A, 7B)**. Notably, *Tmem115* mutants showed a stronger effect than *Tmf* mutants, and the *Tmem115:Tmf* double mutants showed a further decrease of Golgi width, suggesting that Tmem115 has functions beyond localising Tmf.

This idea is supported by observations that in mammalian cells TMEM115 knockdowns caused more changes in the Golgi proteome than TMF1 knockdowns (**Figs. 7C**), and that TMEM115 and TMF1 knockdown Golgi proteomes showed distinct GO-CC features: in a range of selected GO-CC terms, whereas both knockdowns perturb ECM protein transport, TMF1 knockdown resulted in altered Golgi abundance of cytoskeleton and endosomal recycling proteins, and TMEM115 knockdown mostly altered the Golgi abundance of lysosomal, ER, and V-ATPase proteins (**Fig. 7D**).

Finally, we also assessed the phenotype of loss of TMEM115 in mice by making a complete knockout of the gene (**Fig. S13A**). TMEM115 KO mice are born at Mendelian ratios but die postnatally, and the few survivors struggle to gain weight and eventually die before weaning (**Figs. 7E, 7F, S13B, S13C**). The phenotype of TMF1 KO mice has been reported to be much less severe: homozygous knockouts appear healthy (only sperm and metabolic defects have been reported (Lerer-Goldshtein et al., 2010; Rahimi et al., 2021)). Thus, mouse genetics supports the conclusion that TMEM115 has a wider biological function than TMF1, implying that the molecular role in localising TMF1 is not its sole function.

## Discussion

Here we report that the rhomboid-like superfamily member TMEM115 is an essential component of the machinery that localises the golgin TMF1 to its specific position at the Golgi rim. Our discovery was a surprise for two reasons. First, the small GTPase Rab6A is not, as previously reported, sufficient to localise TMF1. Our data highlight the significance of two-factor addressing of TMF1 by the combined action of TMEM115 and Rab6A. In the absence of TMEM115, TMF1 does not move to other Rab6A-specific cellular compartments; instead, it is simply de-localised from the Golgi (**Fig. 7G**). Even when the GTP-locked, constitutively activated form of Rab6A is highly overexpressed, it is still inefficient at localising TMF1, except in the presence of the cytoplasmic C-terminus of TMEM115. Notably, in our relocalisation experiments, Rab6A and Rab5A also both failed to efficiently relocate their well-known effectors BICD2 and EEA1, implying that the inefficiency of Rab6A in localising TMF1 is not an isolated case.

GTPases of the Rab, Arf and Arl family are central to the accuracy of targeting of peripheral membrane proteins and thereby to cellular protein trafficking. Indeed, they represent the textbook explanation of protein targeting. However, the small GTPase-effector model cannot easily explain the observations that some Rabs display partially overlapping organellar localisation and many proteins can recognise more than just one GTP-bound Rab but are not targeted by all of them (Gillingham et al., 2014; Grosshans et al., 2006; Hayes et al., 2009; Sinka et al., 2008; Vitale et al., 1998). Nor does a simple version of the small GTPase-effector model easily account for membrane targeting GTPases having significant levels of similarity, both in structure and sequence, but very distinct effector targeting functions. It is also striking that the affinity between Rabs and their effectors is generally weak and their interactions unstable *in vitro*, but they are nevertheless able to mediate long-range transport *in vivo* (Brault et al., 2022; Grigoriev et al., 2007; Noell et al., 2018).

A coincidence detecting mechanism for increasing the affinity of Rab5-EEA1 binding and endosomal targeting has previously been proposed (Christoforidis et al., 1999; Simonsen et al., 1998). In that case, a local accumulation of membrane lipid PtdIns(3)P (PI3P) was shown to stabilise the Rab5-EEA1 interaction but does not contribute to precision of EEA1 targeting since PI3 kinase, which generates PI3P, is itself localised by Rab5; *i.e.* both elements of the coincidence detector depend on the same Rab. In contrast, we have shown that TMEM115 and Rab6A provide independent localisation information for TMF1. Overall, we suggest that, *in vivo*, cofactors quite commonly provide Rabs, and possibly other membrane targeting GTPases, with additional selectivity and affinity for effectors.

The second surprise in our results is a challenge to the theme that until now has linked all rhomboid-like proteins, both proteases and pseudoproteases: that specific TMD recognition lies at the heart of their action (Freeman, 2014; Tichá et al., 2018). In the case of TMEM115, only the short helical motif in the C-terminal cytoplasmic domain is needed to form the TMEM115-TMF1-Rab6A complex, implying that rhomboid-like proteins can have functions that are independent of TMD recognition. The evolutionary conservation of TMEM115 in TMF1 localisation suggests that this role is ancient. It is important to note, however, that our results do not necessarily mean that the rhomboid-like TMD module of TMEM115 has no function. Both our proteomic and genetic data imply that TMEM115 contributes to mechanisms beyond localising TMF1. These inferred additional functions could involve canonical rhomboid-like TMD recognition.

TMEM115, Rab6A, and TMF1 assemble into a ternary complex in which each of them is essential *in vivo*. Within the complex, two Rab6A molecules bind to the TMF1 coiled-coil dimers, as expected when compared to canonical Rab-effector binding. TMEM115 contributes to the assembly via a short helix in its otherwise unstructured cytoplasmic C-terminal domain. One consequence of the addition of TMEM115 to the complex is spatial: Rab6A has a short tail that links its compact globular GTPase domain to its lipid anchor, while the link between the interacting helix in TMEM115 and the TM core that tethers it to the membrane is longer and more flexible. We speculate that the extended reach of TMEM115 might allow it to establish contact with TMF1 on incoming vesicles before Rab6A can.

Our discovery of TMEM115 as an essential cofactor in Rab6A-dependent localisation of TMF1 does not rule out the existence of further new components of the addressing machinery. Indeed, a possible explanation for our difficulty in identifying a stable direct interface between TMEM115 and Rab6A, despite their interdependency for TTR complex formation *in vivo*, would be if other factors contribute to the TMEM115-Rab6A interaction. Furthermore, the unexpected lack of an identified function for the main rhomboid-like membrane domain of TMEM115, as well as the observations that Rab5A is inefficient at localising its effector EEA1, are both consistent with the involvement of unknown cofactors. Overall, we propose that the role for TMEM115 in Rab6-dependent protein localisation may exemplify an underappreciated network of membrane targeting cofactors, working in concert with small GTPases to ensure the correct addressing of membrane and secreted proteins.

## Materials and Methods

### Cell-lines, Crispr/Cas9-mediated gene KO, and generation of stable cells

hTERT-RPE1, HEK-293T, COS7, Hela, MCF7, and MEF cells were cultured in DMEM/F12 (Gibco, Cat. #D8437) or DMEM high glucose medium (Gibco, Cat. #D6429), supplemented with 10% foetal bovine serum (Gibco, Cat. #F4135), and Penicillin and Streptomycin (1:1000 dilution, Gibco, Cat. #15140-122), in a 37°C, 5% CO2 incubator. Cell passage was conducted following ATCC’s cell culture instructions and cells are regularly tested to be mycoplasma-free.

To establish TMEM115 KO RPE1 cells, two guide RNAs (gRNA) were used to cut out part of TMEM115 (5’-GACGGTGAAGCGCTACGATG-3’ and 5’-GACTTATTAAGTGGCCCGAG-3’). 600,000 cells were spun down, washed with PBS and resuspended in 30 μL of Buffer R (Neon nucleofection system). 1 μL of gRNA each was mixed with 1 μL of Tracer RNA, incubated for 5 min at 95°C, cooled to room temperature (RT), then mixed with 0.75 μL of Alt-R S.p. HiFi Cas9 Nuclease V3 (IDT DNA, Cat. #1081060) and incubated for 20 min at RT. 10 μL of cells were electroporated with Neon Nucleofector (Thermo) using the program 1350V, 10 ms, 3 pulses. 1.5 μL from each gRNA-tracer-Cas9 mix were mixed in a new tube and 20 μL of RPE1 cells were added and mixed. 10 μL was electroporated at 1700V, 30 ms, 1 pulse. After nucleofection, the cells were seeded in full medium for recovery for 3 days. Some cells were collected for a PCR genotyping using the primers: 5’- CTGACCACTTTGCTTTCGCC-3’ and 5’-GAAATTGGAGTGGCCCAGA-3’. Single clonal cells were sorted by FACS and were verified again by genomic sequencing and western blot.

To create stable RPE1 cells, human TMEM115-3xHA was first cloned into the MSCV-IRES-eGFP lentivirus vector, followed by site-directed mutagenesis or truncation. DNAs were mixed with the virus packaging plasmids pMD2.G and pSPAX2 in a 10:3:7 ratio and incubated with FuGENE HD (Promega, Cat. #E2312) in a 1:4 ratio in OptiMEM (Gibco, Cat. #31985) for 15 min at RT. HEK-293T cells were transfected with the DNA-FuGENE mixtures for 1 day, changed with fresh medium. Three days later, supernatants were collected and filtered through 0.45 μm filters and added into the RPE1 WT or TMEM115 KO6 cells, in the presence of 10 μg/mL polybrene (Sigma, Cat. #TR-1003) for 1 day. Positive cells were sorted by fluorescent FACS.

### Mouse genetics and viability analysis

All procedures on mice were conducted in accordance with the UK Scientific Procedures Act (1986) under a project license (PPL) authorized by the UK Home Office Animal Procedures Committee and approved by the Sir William Dunn School of Pathology Local Ethical Review Committee. Mice were maintained under specified pathogen free conditions in accredited facilities at the University of Oxford, and were kept at constant temperature and humidity, at a 12h/12h light/dark cycle and had ad libitum access to food and water. *Tmem115^+/-^*mice were obtained as indicated in the section below. Given the deleterious phenotype of the *Tmem115^-/-^* mice, all litters produced from crossing of two heterozygous parentals were genotyped after birth, monitored, and weighed daily. According to Project Licence PP5666180, when an animal showed a loss of 15% of its peak weight, it was humanely culled by a Schedule 1 Method. Matching WT and heterozygous littermates were also culled for comparison.

*Tmem115* mutant mice were generated using embryonic stem cells from the Knockout Mouse Project (KOMP) repository in which a 3596-bp sequence, including all translated sequence of exon 1 and most of exon 2, had been replaced with a lacZ reporter and a neomycin resistance cassette (Project VG10466, **Fig. S13A**). These cells were injected into C57Bl/6J blastocysts to generate chimeric animals from which the first heterozygous mice were obtained. Heterozygous mice were inter-crossed to produce *Tmem115^+/+^* (WT), *Tmem115^+/-^* (heterozygous, HT) and *Tmem115^-/-^* (KO) animals. All animals coming from these crosses were monitored daily and weighed.

Genotyping was performed using a combination of the three following primers 5’- CGGAATCCAGTCTCATCAC-3’ (WT-specific), 5’-CAGCAGCCTCTGTTCCACA-3’ (KO-specific) and the common reverse primer 5’-GGCATTGTGGTTGTCTGGA-3’.

### Fly genetics and immuno-fluorescence microscopy of fly tissues

#### Drosophila stocks

All fly stocks were kept at 25°C using a 12h-light-12h-dark cycle. Fly stocks lacking CG9536 were generated using the FLP-FRT deletion strategy established by Exelixis (Parks et al., 2004; Thibault et al., 2004). piggyBac elements pBacRBe02233 and pBacRBe04259 were recombined to delete the entire coding sequence of CG9536 (**Fig. S13D**). The precise recombination sites were confirmed by genomic sequencing. Two genetically identical mutant alleles, *CG9536*^3^ and *CG9536*^28^, were produced in this way. The *TmfΔ19* and GFP- Tmf strains were described in a previous study (Park et al., 2022). Female flies with two copies of GFP-Tmf were used to generate the GFP-Tmf; *CG9536^3^*mutants, and the *TmfΔ19;CG9536^3^* flies were generated by crossing the two individual mutant strains. The Oregon-R strain was used as the WT control.

#### Imaging 3rd instar larval salivary glands and analyzing GFP-Tmf protein levels

To analyze the role of CG9536 in regulating Golgi-associated GFP-Tmf levels, the salivary glands (SGs) of GFP-Tmf;*CG9536^3^* mutant flies and GFP-Tmf control flies from wandering 3rd instar larvae were dissected in Schneider’s medium and fixed in 4% PFA for 30 min at 25°C. SGs were washed in 0.1% Triton X-100 PBS, twice, 15 min each, then equilibrated in Vectashield containing DAPI (1 μg/ml, Vector Laboratories) overnight at -20°C. SGs were mounted in Vectashield containing DAPI and images were taken on a Zeiss LSM 900 with Airy Scan 2. Three repeats were carried out for each genotype. To measure Golgi-associated GFP-Tmf intensities, the same settings were used for each genotype and all technical repeats. The mean intensity was measured in FIJI using a macro to set the same threshold for all images. The number of detected Golgi per image was roughly the same for control and mutants.

To analyse the total GFP-Tmf protein levels, five SGs were dissected from wandering 3rd instar larvae in Schneider’s medium for each genotype. SGs were immediately transferred into 20 µL of RIPA buffer containing protease inhibitors (Roche, Cat. # 05892970001). SGs were homogenised and left on ice for 25 min. NuPAGE 4 x LDS sample buffer (Invitrogen, Cat. #NP0007) with 5% β-mercaptoethanol was added and samples were boiled at 90°C for 10 min. Samples were separated on a 4-12% Bis-Tris gel (Invitrogen), followed by an ordinary WB procedure. Mouse anti-GFP (Roche, Cat. #11814460001) and mouse anti-Actin (Sigma, Cat. #A4700) were used.

#### Golgi width measurement

For Golgi width analysis, 3rd instar larval SGs were dissected in Schneider’s medium and fixed in 4% PFA for 30 min at 25°C. SGs were first permeabilised in 0.3% Triton X-100 PBS for four times, 30 min each, then blocked with 0.1% Triton X-100 plus 5% BSA PBS for four times, 30 min each. Primary antibodies were incubated in 0.1% Triton X-100 plus 5% BSA PBS overnight at 4°C. SGs were washed in 0.1% Triton X-100 PBS for four times, 30 min each, then incubated with secondary antibodies in 0.1% Triton X-100 plus 5% BSA PBS overnight at 4°C. SGs were washed in 0.1% Triton X-100 PBS for four times, 30 min each, then equilibrated in Vectashield containing DAPI (1 μg/mL, Vector Laboratories) overnight at -20°C. SGs were imaged on a Zeiss LSM 900 with Airy Scan 2. Rabbit anti-GM130 (Abcam, Cat. #ab30637), mouse anti-Golgin84 and goat anti-Golgin245 (Riedel et al., 2016) were used. Images were taken from 2-3 different larvae per experiment, three technical repeats. To measure Golgi width from GM130 confocal micrographs, lines were drawn across the widest point to measure the width of the stack. At least 30 Golgi stacks were measured per image.

### Transient transfection and siRNA-mediated gene knockdown

For transient transfection, plasmid DNAs were mixed with FuGENE in a 1:3 ratio and incubated in OptiMEM for 15 min at RT. Cells were changed to antibiotic-free full medium during transfection. Five hours later, cells were changed to full medium, and 24 h later were fixed for imaging or lysed for GST-pulldown. In the case of the mitochondria anchoring-away experiments, RPE1 cells were fixed at 72 h post transfection.

To knock down the genes of interest, RPE1 cells in a 35 mm dish were at 60% confluency on the day of transfection. 30 pmol siRNAs (Ambion) were mixed with 3 μL of Lipofectamine RNAiMax (Invitrogen, Cat. #13778150) in OptiMEM and incubated for 15 min at RT. The mixture was added to the cells. On the following day, the cells were passaged or seeded on coverslips and cultured for another 3 days. The following siRNAs were used: for TMEM115, 5’-UGGUCUACCUGUUCACUGUtt-3’ (Ambion, s21819) and 5’- CGGCGGUACUAUUCCUCUAtt-3’ (Ambion, s21817); for Rab6A/6A’, 5’- GAGCUGAAUGUUAUGUUUAtt-3’ (Ambion, s11685) and 5’- AAUUCUUCACGGUAAGAAAtt-3’ (Ambion, s11686); for TMF1, 5’- GACAUAGCUUUGGAACCUAtt-3’ (Ambion, s14228) and 5’- CUAAGAGAUUUGGAUCAAAtt-3’ (Ambion, s14227); negative control (Ambion, Cat. #4390847).

### Immuno-precipitation and Western blot

Immuno-precipitation (IP) and Western blot (WB) were carried out as previously described (Zhang et al., 2019) with modifications. Briefly, cell pellets were lysed with the IP buffer (1% Digitonin, 150 mM NaCl, 50 mM HEPES, pH 7.4) for IP and GST pulldown assays or the RIPA buffer (1% NP40, 0.5% Sodium Deoxycholate, 50 mM NaCl, 50 mM Tris-HCl pH 7.4) for WB, both supplemented with cOmplete Protease inhibitors (Roche, Cat. #81410500).

The supernatants were incubated with anti-HA (Pierce, Cat. #88837, or Cell Signalling, Cat. #11846), anti-Flag (Pierce, Cat. #A36797), or anti-GFP (Cell Signalling, Cat. #67090, or Proteintech, Cat. #gta-100) beads at 4°C for 1.5 h. For GST-pulldown experiments, 6 μg of purified GST and GST-tagged proteins were diluted in 1% Triton in PBS and re-coated onto the Glutathione magnetic beads (Pierce, Cat. #G0924) at 4°C for 1.5 h. Cells were lysed with the IP buffer, and the supernatants were added into the already coated beads and incubated at 4°C for 1.5 h.

After incubation, beads were washed with IP buffer three times, 10 min each at 4°C. Samples were heated at 65°C for 7 min, then stored at -20°C or processed immediately for WB. Antibodies used for WB were anti-TMEM115 (Atlas, Cat. #HPA015497), TMF1 (Atlas, Cat. #HPA008729), Rab6A/6A’ (Novus, Cat. #NBP1-33110), Rab33B (GeneTex, Cat. #GTX116390), GM130 (Proteintech, Cat. #11308-1-ap), GMAP210 (Proteintech, Cat. #26456-1-ap), Golgin160 (Proteintech, Cat. #21193-1-ap), GCC1 (Proteintech, 16271-1-ap), Golgin97 (Proteintech, Cat. #12640-1-ap), BICD2 (Atlas, Cat. #HPA023013), ADAMTS12 (Proteintech, Cat. #24934-1-ap), COL4A1 (Proteintech, Cat. #30850-1-ap), SNCA (Proteintech, Cat. #10842-1-ap), CTSS (Cell Signal Tech, Cat. #25084), HSP90 (Proteintech, Cat. #17260-1-ap), GAPDH (Proteintech, Cat. #60004-1-Ig), HA tag (Cell Signaling, Cat. #3724), Flag tag (Proteintech, Cat. #20543-1-ap).

### Immuno-fluorescence microscopy of mammalian cells and data processing

Imaging experiments were carried out as previously described (Zhang et al., 2019). Briefly, cells were fixed with 4% PFA PBS for 20 min at RT and permeabilized with 0.2% Triton X- 100 PBS for 5 min at RT. Cells were incubated with the indicated primary antibodies overnight at 4°C overnight and washed with PBS and incubated with corresponding secondary antibodies for 2 h at RT. DNA was stained with DAPI. Images were captured using the Carl Zeiss LSM880 fluorescence microscope with the confocal mode or the Airyscan super-resolution mode. Fluorescence intensity of the indicated proteins was quantified using FIJI by manually drawing a polygonal border around the areas of interest, and the mean intensity of the areas was used for downstream analysis.

Critical antibodies involved in imaging were used in a 1:200 dilution: anti-TMEM115 (Atlas, Cat. #HPA015497), TMF1 (Atlas, Cat. #HPA008729), Rab6GTP (AdipoGen, Cat. #AG-27B- 0004), GM130 (BD, Cat. # 610823), Giantin (BioLegend, Cat. #924301), GMAP210 (Proteintech, Cat. #26456-1-ap), Golgin160 (Proteintech, Cat. #21193-1-ap), GCC1 (Proteintech, 16271-1-ap), Golgin97 (Proteintech, Cat. #12640-1-ap), TGN46 (GeneTex, Cat. #GTX74290), EEA1 (BD, Cat. #610456), BICD2 (Atlas, Cat. #HPA023013), HA tag (Invitrogen, Cat. #26183), Flag tag (Proteintech, Cat. #20543-1-ap, 66008-3-lg), Myc tag (Proteintech, Cat. #16286-1-ap). Fluorescent secondary antibodies were from Invitrogen (Cat. #A-11008, A-11001, A-11011, A-11004, A-31573, A-31571) and Abcam (Cat. #ab150179).

### Protein expression and purification

#### Purification of full-length Rab6A and the TMF1 and TMEM115 truncates for Cryo-EM

Full-length human Rab6A, human TMF1 (residues 972-1093), human TMEM115 (residues 275-305) were expressed in E. coli strain BL21 (DE3) from a pET28a vector modified to encode an N-terminal His6-SUMO fusion, or in the case of TMEM115, a pLIP vector which encodes two TEV protease-cleavable, His-tagged lipoyl domains fused at either terminus of the insert. Protein expression was induced by using 1 mM Isopropyl ß-D-1- thiogalactopyranoside (IPTG) for 12 h at 18°C, or in the case of TMEM115, 1 mM IPTG for 3 h at 37°C. Cells pellets were resuspended in Buffer A (50 mM Tris-HCl pH 8.0, 500 mM NaCl, 1 mM EDTA, and 2 mM imidazole) supplemented with DNase I (20 μg/mL), lysozyme (500 μg/mL) and cOmplete EDTA-free Protease Inhibitor Cocktail (Sigma) for 30 min before passing through an EmulsiFlex C5 homogenisor (Avestin) at 15,000 psi. Lysates were clarified by centrifugation (40,000xg, 40 min), incubated with 5 mL of cOmplete His-tag purification resin (Roche), and then the resin was washed with 40 column volume (CV) of buffer A.

Proteins were eluted in 6 CV of buffer B (50 mM Tris-HCl pH 8.0, 150 mM NaCl, 1 mM EDTA, and 200 mM imidazole). The eluate was supplemented with 1 mg of Saccharomyces cerevisiae Ubiquitin-like-specific protease 1 (Ulp1), or in the case of TMEM115, 5 mg of TEV protease, and dialysed for 12 h into buffer C (20 mM HEPES pH 7.4, 150 mM NaCl, 1 mM EDTA) using 10 kDa molecular weight cutoff (MWCO) SnakeSkin dialysis tubing (Thermo Scientific). The dialysed material was incubated with 5 mL of cOmplete His-tag purification resin (Roche) for 1 h and the resin flow through loaded onto an S200 16/600, or in the case of TMEM115, S75 16/600, size-exclusion chromatography column (Cytiva), pre-equilibrated with buffer D (20 mM HEPES pH 7.4, 150 mM NaCl). Peak fractions were collected and concentrated using a 10 kDa, or in the case of TMEM115, 3 kDa, MWCO Vivaspin (GE healthcare) centrifugal filter unit. Proteins aliquots were stored at -80°C.

#### Purification of full-length TMEM115 for cryo-EM

Full-length human TMEM115 fused to a mVenus-twin-Strep tag fusion was produced in HEK293S GnTI- cells stably expressing protein under the control of a tetracycline operator. Cells were grown in suspension in Freestyle media to a cell density of 3 x 10^6^ at 37°C, expression was induced by adding 10 μg/mL doxycycline and incubated for a further 48 h at 37°C. Cell pellets were resuspended in PBS supplemented with DNase I (20 μg/mL) and a cOmplete EDTA-free Protease Inhibitor Cocktail tablet (Sigma) for 30 min before passing through an EmulsiFlex C5 homogenisor (Avestin) at 15,000 psi. Lysates were clarified by centrifugation (12,000xg, 20 min) and membranes were harvested by ultracentrifugation (150,000xg, 90 min). Membranes were resuspended in buffer E (50 mM sodium phosphate pH 8.0, 150 mM NaCl) and solubilized by incubation with 1% (w/v) Lauryl maltose neopentyl glycol (LMNG; Anatrace) for 1 h. Insoluble material was removed by centrifugation (100,000xg, 30 min) and solubilized membranes applied to a 5 mL Strep-Tactin XT 4Flow column (IBA). The column was washed with 10 CV of buffer E containing 0.02% (w/v) LMNG. Proteins were eluted in 4 CV of buffer E containing 0.02% (w/v) LMNG and 50 mM biotin.

#### Purification of GST-TMF1 fragments

Mouse TMF1 fragments and mutants were expressed in the E. coli strain BL21 (DE3) in the pGEX-4T-1 vector. Protein expression was induced by using 1 mM IPTG for 12 h at 18°C. Cells pellets were washed with PBS, then lysed in BugBuster (Millipore, Cat. #70584) supplemented with cOmplete EDTA-free Protease Inhibitor Cocktail, for 15 min on ice. The supernatants were clarified by centrifuging at 5000xg and then at 20, 000xg for 15 min each at 4°C. The lysates were incubated with the Glutathione Sepharose 4B beads (Cytiva, Cat. #17075601) for 1.5 h with rotation at 4°C, washed with 30 CV of cold PBS, then eluted with buffer (20 mM reduced glutathione, 1 mM DTT, 200 mM NaCl, 200 mM Tris-HCl pH 8.0). Eluate was concentrated by centrifuging with the 10 kDa or 3 kDa MWCO filter units (Amicon, Cat. #UFC9010, UFC9030) depending on the size of target proteins. Proteins aliquots were stored at -80°C.

### Size-exclusion chromatography analysis of protein complex assembly

Full-length Rab6, TMF1 (residues 972-1093) and/or TM115 (residues 275-305) were mixed in a 2:1:2 molar ratio in the presence of 1 mM MgCl2 and 1 mM GTPγS. The mixture was incubated for 12 h on ice, then loaded onto a S200 10/300 size-exclusion column pre-equilibrated with buffer (20 mM HEPES pH 7.4, 150 mM NaCl, 1 mM MgCl2). Eluate fractions were analysed by SDS-PAGE and Coomassie staining.

### Cryo-electron microscopy and data processing

To assemble the TMEM115-TMF1 (residues 972-1093)-Rab6A complex, purified TMF1 (0.5 mL at 3.3 mg/mL) and Rab6A (0.5 mL at 2.65 mg/mL) were added to the full-length TMEM115 eluate along with 100 μg of His-tagged TEV protease (to remove the C-terminal mVenus tag from full-length TMEM115) and the mixture was incubated for 12 h at 4°C. The sample was incubated with 1 mL of cOmplete His-tag purification resin for 1 h and the resin flow was concentrated to 0.5 mL. The sample was loaded onto an S6 10/300 size-exclusion chromatography column pre-equilibrated in buffer (150 mM NaCl, 0.02% LMNG, 20 mM HEPES pH 7.4). Eluate fractions were analysed by SDS-PAGE and Coomassie staining.

To assemble the TMEM115 (residues 275-305)-TMF1 (residues 972-1093)-Rab6A complex, Rab6A (200 μL at 241.6 μM), TMF1 (100 μL at 227 μM), and TMEM115 (200 μL at 249.6 μM) were mixed with 0.5 μL MgCl2 (1 M stock), 5 μL GTPγS (100 mM stock), and 50 μL of buffer D and incubated for 12 h on ice. This mixture was loaded onto a S200 10/300 size-exclusion column pre-equilibrated with buffer E (20 mM HEPES pH 7.4, 150 mM NaCl, 1 mM MgCl2). Peak fractions corresponding to the complex were collected and concentrated using the 10 kDa MWCO Vivaspin (GE healthcare) filter unit to A280nm = 0.8 to prepare grids without additives or concentrated to A280nm = 8.2 for samples supplemented with 0.7 mM fluorinated octyl maltoside (Anatrace).

Samples (4 μL each) were adsorbed to glow discharged holey carbon-coated grids (Quantifoil 300 mesh, Au R1.2/1,3) for 30 s at 15 mA. Grids were then blotted for 2 s at 100% humidity at 4°C and frozen in liquid ethane using a Vitrobot Mark IV (FEI). Data were collected on the Thermo Scientific CFEG-equipped Titan Krios G4 microscope, operating at 300 kV with a Selectris X imaging filter with slit width of 10 eV at 165,000x magnification on a Falcon 4 direct detection camera. Pixel size was 0.693 Å and movies were collected at a total dose ranging from 54.0 e-/A2 – 61.9 e-/A2. Patched motion correction, CTF parameter estimation, particle picking, extraction, and initial 2D classification were performed in SIMPLE 3.0. All downstream processing was carried out in cryoSPARC, using the csparc2star.py script within UCSF pyem to convert between formats.

### Immuno-transmission electron microscopy

Stable RPE1 TMEM115-HA Cells were collected and washed with PBS, resuspended in 2% PFA and 0.1% glutaraldehyde in PIPES buffer (0.1 M, pH 7.4), then incubated for 2 h at RT ahead of storage at 4°C overnight. Samples were washed in PIPES buffer and infiltrated in 12% pork gelatine for 5 min at 37°C, centrifuged to form a loose pellet, then incubated for 20 min at 4°C to allow gelatine to solidify. The gelatine-embedded samples were dissected into small blocks and infiltrated in 2.3 M sucrose at 4°C overnight with rotation. Selected blocks were mounted on cryo-ultramicrotomy pins and snap-frozen in liquid nitrogen. Ultrathin cryosections (∼100 nm) were collected and mounted on 50 mesh copper TEM grids coated with formvar and carbon.

For immunogold labelling, sections were incubated in an excess of PBS for 30 min at 37°C, then rinsed for 2 min, 3 times, in droplets of PBS with 20 mM glycine. Sections were blocked for 15 min in blocking buffer (0.1% Aurion BSA-c, 0.5% cold water fish skin gelatine in PBS). Sections were incubated with the anti-HA antibody (Cell Signaling, Cat. #3724, 1:20 dilution) for 90 min in blocking buffer at RT. Sections were washed for 2 min, 6 times, in blocking buffer, then incubated with secondary antibody (CMC Utrecht Protein-A-gold, 10nm, 1:20 dilution) in blocking buffer for 45 min at RT. Sections were washed 2 min, 5 times, in PBS, fixed for another 5 min in 1% glutaraldehyde in PBS then washed again in milliQ water for 1 min, 6 times. Sections were stained with a mixture of 2% methylcellulose and 2% uranyl acetate (in 85:15 ratio) for 10 min on ice. Grids were looped out of the staining solution, excess solution was side-blotted away and the grids were allowed to air dry. Samples were imaged using a JEOL Flash 120kV TEM equipped with a Gatan Rio camera.

### Isolation of Golgi membranes and total vesicle fractions

Golgi membranes were isolated by using an ultracentrifugation and sucrose gradient-based methodology (Graham, 2001) with necessary adjustments. Briefly, RPE1 cells from eight 15 cm dishes at a confluency of 80% were gently scraped down in PBS. The cell pellets were resuspended in 1.8 mL HM buffer (0.25 M sucrose, 10 mM Tris-HCl pH 7.4) supplemented with cOmplete Protease inhibitors and transferred into a ball-bearing homogenizer (Kimble, Cat. #885301). The cell suspension was homogenized with the B-type pestle for 26 strokes, then made into 1.4 M sucrose homogenate by adding 2 volumes of the sucrose stock (2.0 M sucrose, 10 mM Tris-HCl pH 7.4). The homogenate was then transferred directly to the bottom of an ultracentrifugation tube (Beckman, Cat. #344059) and extremely gently sequentially layered with 4.5 mL of 1.2 M sucrose and 2 mL of 0.8 M sucrose in 10 mM Tris-HCl. The tubes were centrifuged at 110,000xg, 4°C for 2 h, with acceleration speed “5” and “coast” to stop. After centrifugation, the Golgi membranes were gently collected from the 0.8 M / 1.2 M sucrose interface and transferred into new tubes (Beckman, Cat. #344088). The fractions were diluted with 2 volumes of PBS, and further centrifuged at 100,000xg, 4°C, for 26 min. The pellets were gently washed with 2 mL PBS and then lysed with the RIPA buffer. Total protein concentration was measured using the BCA method (Pierce, Cat. #23227).

Total vesicles were isolated following an ultracentrifugation and iodixanol gradient-based methodology (Zhang et al., 2020) with modifications. Briefly, RPE1 cells from one 15 cm dish at a confluency of 80% were harvested and lysed in 1 mL of HB1 buffer (400 mM sucrose, 1 mM EDTA, 0.3 mM DTT, 20 mM HEPES-KOH pH 7.2) supplemented with protease inhibitors by quickly passing through a 23G needle for 26 times. The lysate was centrifuged at 1,000xg for 10 min and the supernatant was centrifuged at 20,000xg for 30 min. The supernatant was then transferred to a new tube (Beckman, Cat. #344088) and centrifuged at 100,000xg for 1 h. The transparent membrane pellets were resuspended with 250 μL of 35% Optiprep (Sigma, Cat. #D1556, diluted from 60% stock with B88 buffer (250 mM sorbitol, 150 mM KAc, 5mM MgAC2, 20mM HEPES pH 7.2)). The 35% fraction was transferred to a new tube (Beckman, Cat. #347357), gently sequentially layered with 700 μL of 30% Optiprep diluted in B88 buffer and 120 μL of B88 buffer, then centrifuged at 150,000xg, 4°C, for 2.5 h. The membrane fractions of 100 μL each were sequentially collected from the top, stored at -20°C or immediately processed for WB. Vesicles were primarily enriched in the second 100 μL sample from the top.

### Proteomics analysis and data processing

#### TMT Labelling and High pH reversed-phase chromatography

Aliquots of 30 µg of Golgi membrane extract were digested with trypsin overnight, labelled with Tandem Mass Tag (TMTpro) sixteen-plex reagents according to the manufacturer’s protocol (Thermo Scientific) and the labelled samples were pooled. An aliquot of 200 μg of the pooled sample was desalted using a SepPak cartridge according to the manufacturer’s instructions (Waters). Eluate from the SepPak cartridge was evaporated to dryness and resuspended in buffer A (20 mM ammonium hydroxide, pH 10) prior to fractionation by high pH reversed-phase chromatography using an Ultimate 3000 liquid chromatography system (Thermo Scientific). In brief, the sample was loaded onto an XBridge BEH C18 Column (130Å, 3.5 µm, 2.1 mm X 150 mm, Waters) in buffer A and peptides were eluted with an increasing gradient of buffer B (20 mM Ammonium Hydroxide in acetonitrile, pH 10) from 0- 95% over 60 min. The resulting fractions (concatenated into 15 in total) were evaporated to dryness and resuspended in 1% formic acid prior to analysis by nano-LC MS using an Orbitrap Fusion Lumos mass spectrometer (Thermo Scientific).

#### Nano-LC Mass Spectrometry

High pH RP fractions were further fractionated using an Ultimate 3000 nano-LC system in line with an Orbitrap Fusion Lumos mass spectrometer (Thermo Scientific). In brief, peptides in 1% (vol/vol) formic acid were injected onto an Acclaim PepMap C18 nano-trap column (Thermo Scientific). After washing with 0.5% acetonitrile in 0.1% formic acid, peptides were resolved on a 500 mm × 75 μm Acclaim PepMap C18 reverse phase analytical column (Thermo Scientific) over a 150 min organic gradient, using 7 gradient segments (1-6% solvent B over 1min, 6-15% solvent B over 58min, 15-32% solvent B over 58min, 32-40% solvent B over 5min, 40-90% solvent B over 1min, held at 90% solvent B for 6 min and then reduced to 1% solvent B over 1min) with a flow rate of 300 nL/min. Solvent A was 0.1% formic acid, and Solvent B was aqueous 80% acetonitrile in 0.1% formic acid. Peptides were ionized by nano-electrospray ionization at 2.0 kV using a stainless-steel emitter with an internal diameter of 30 μm (Thermo Scientific) and a capillary temperature of 300°C.

All spectra were acquired using an Orbitrap Fusion Lumos mass spectrometer controlled by Xcalibur 3.0 software (Thermo Scientific) and operated in data-dependent acquisition mode using an SPS-MS3 workflow. FTMS1 spectra were collected at a resolution of 120,000, with an automatic gain control (AGC) target of 200,000 and a max injection time of 50 ms. Precursors were filtered with an intensity threshold of 5,000, according to charge state and with monoisotopic peak determination set to Peptide. Previously interrogated precursors were excluded using a dynamic window (60 s ± 10 ppm). The MS2 precursors were isolated with a quadrupole isolation window of 0.7 m/z. ITMS2 spectra were collected with an AGC target of 10,000, max injection time of 70 ms and CID collision energy of 35%.

For FTMS3 analysis, the Orbitrap was operated at 50,000-resolution with an AGC target of 50,000 and a max injection time of 105 ms. Precursors were fragmented by high energy collision dissociation (HCD) at a normalized collision energy of 60% to ensure maximal TMT reporter ion yield. Synchronous Precursor Selection (SPS) was enabled to include up to 10 MS2 fragment ions in the FTMS3 scan.

#### Protein identification and quantification

The raw data files were processed and quantified using Proteome Discoverer software v2.4 (Thermo Scientific) and searched against the UniProt Human database (downloaded Jan 2024: 82415 entries) using the SEQUEST HT algorithm. Peptide precursor mass tolerance was set at 10 ppm, and MS/MS tolerance was set at 0.6 Da. Search criteria included oxidation of methionine (+15.995 Da), acetylation of the protein N-terminus (+42.011 Da) and Methionine loss plus acetylation of the protein N-terminus (-89.03 Da) as variable modifications and carbamidomethylation of cysteine (+57.0214) and the addition of the TMTpro mass tag (+304.207) to peptide N-termini and lysine as fixed modifications. Searches were performed with full tryptic digestion and a maximum of two missed cleavages were allowed. The reverse database search option was enabled, and all data was filtered to satisfy false discovery rate (FDR) of 5%.

Protein groupings were determined by PD2.4, however, further the master protein selection was improved, and all following statistical analyses were performed in the R statistical environment version 4.4.0. The Master protein improvement script first searches Uniprot for the status of all protein accessions and updates redirected or obsolete accessions. The script further takes the candidate master proteins for each group and uses current Uniprot review and annotation status to select the best annotated protein as master protein without loss of identification or quantification quality.

The protein abundances for each sample were normalized using a sum of peptide abundance before being Log2-transformed. The data were statistically analysed using normalized abundances. A single LIMMA model was constructed for all conditions and moderated t-tests performed for each comparison of interest, then the p-value was adjusted using the Benjamini-Hochberg FDR method. Outputs for raw abundance were exported to Excel and tidied up for ease of use. For each comparison, using both raw and normalized data, the -log10 (*P*-value) of each protein was plotted against the log2 (fold change).

### Protein structure prediction, topology and sequence alliance analysis

Formation of the TTR complex was predicted by using AlphaFold 3 (https://alphafoldserver.com/) and the interacting residues were analysed by using MAPIA (https://mapiya.lcbio.pl/). Default algorithms were applied to both programs. Protein hydrophobicity analysis was carried out by using DeepTMHMM (Hallgren et al., 2022) (https://dtu.biolib.com/DeepTMHMM/). Note that the hydrophobicity of TMEM115 can be similarly predicted by many other tools, with only tiny differences in the boundaries. Protein sequence conservation analysis was carried out by using Clustal Omega (https://www.ebi.ac.uk/jdispatcher/msa/clustalo) with the Uniprot IDs of human TMEM115 (Q12893), TMF1 (P82094), Rab6A (P20340); mouse TMEM115 (Q9WUH1), TMF1 (B9EKI3), Rab6A (P35279); fly CG9536 (Q9VMD2), Tmf (Q9W3V2), Rab6 (O18334); yeast Yol107w (Q12239), Sgm1 (P47166), Ypt6 (Q99260).

### Data visualisation

To generate visualised results, the protein quantification data and *P* values were uploaded to SRplot (Tang et al., 2023) (http://www.bioinformatics.com.cn/SRplot) for online processing and graphing. For the co-relation analysis, the protein abundance data and *P* values were uploaded to ProHitsViz (Knight et al., 2017) (https://prohits-viz.org/analysis) for online processing and graphing.

### Statistics

The fluorescent intensity of areas of interest was individually measured in FIJI. For Golgi intensity of the indicated proteins in mammalian cell cultures, at least 30 sample pairs were collected for each group. Statistical analysis was carried out by using unpaired, two-tailed, Students’ t-test, or Kruskal-Wallis test for Figs. 5B, 5C, and 7B.

## Acknowledgements

We thank Drs Kate Heesom and Phil Lewis (from University of Bristol) for MS analysis; Dr Errin Johnson (now at the University of Sydney) and Dr Charlotte Melia (both from the Dunn School EM facility) for TEM analysis; Dr Joey Riepsaame (from Genome Editing Oxford) for gene editing; Dr Jérôme Boulanger (from MRC Laboratory of Molecular Biology, Cambridge) for fly imaging analysis; Professor Daniel Ungar (from University of York) for the mouse TMF1 cDNA, and all other members of the Freeman lab for helpful comments. MF received funding from Wellcome (101035/Z/13/Z and 220887/Z/20/Z), SM received funding from UKRI (United Kingdom Research and Innovation, MC_U105178783), SML received funding from ALSAC and the Intramural Research Program of the NIH. The contributions of the NIH author(s) were made as part of their official duties as NIH federal employees, are in compliance with agency policy requirements, and are considered Works of the United States Government. However, the findings and conclusions presented in this paper are those of the author(s) and do not necessarily reflect the views of the NIH or the U.S. Department of Health and Human Services.

