## Supplemental figures and legends for "The rhomboid-like pseudoprotease TMEM115 defines a two-factor mechanism for Rab6A effector recruitment"

### Supplementary Figures and Legends

**Figure S1 TMEM115 contains six TMDs**

(A) Principle of the TMEM115 topology analysis / antibody accessibility test. 0.2% Triton permeabilises all cellular membranes, whereas 0.2% digitonin only permeabilises the plasma membrane (PM).

(B) TMEM115 contains six TMDs. TMEM115 was tagged at the indicated regions and transiently expressed in RPE1 cells. Under Triton or digitonin permeabilisation, the accessibility of antibodies against the tags were compared. A model of each TMEM115 construct is shown to the left. L, lumen; C, cytoplasm.

Throughout this study, unless otherwise indicated, imaging experiments were done in RPE1 cells and scale bars indicate 10μm; quantification data were derived from at least 30 cells in each group, shown as box-and-whisker plot plus the mean and *P* values.

**Figure S2 TMEM115 is enriched in the Golgi rims and is required for the Golgi localisation of TMF1**

(A) TMEM115 is enriched in the Golgi rims of COS7, Hela, MCF7, and MEF cells, shown by the Airyscan super-resolution microscopy. The boxed areas in the left panels are shown at higher magnification to the right. Giantin indicates the Golgi rim.

(B) Correlation analysis of the two TMEM115 KD Golgi proteomes by the s21819 and s21817 siRNAs. Only consistently altered protein hits across the two KD datasets are plotted.

(C - E) The fly TMEM115 homologous gene, *CG9536*, is required for the Golgi localisation of GFP-Tmf in fly salivary glands. (C) *CG9536* was knocked out in flies expressing endogenously GFP-tagged Tmf. L3 salivary gland cells of the parental and *CG9536*$^{3}$ mutant flies were imaged. The intensities of all Golgi GFP-Tmf per image were quantified and averaged and are shown in (D). (E) The total protein level of GFP-Tmf was not affected in the *CG9536* mutants. L3 salivary gland cells of the parental and *CG9536*$^{3}$ mutant flies were lysed.

**Figure S3 TMEM115 and Rab6A regulate the same pool of Golgi TMF1 and TMEM115 does not regulate the protein level and activity of Rab6A**

(A - E) TMEM115 and Rab6A regulate the same pool of Golgi TMF1. (A) WT and $WT^{GFP}$ cells were singly or doubly knocked down for TMEM115 and Rab6A. (B, D) The indicated cells were co-cultured and immuno-stained for TMF1. (C, E) The effect of non-targeting siRNA (siCon) and TMEM115 and Rab6A KDs on TMF1 Golgi localisation was quantified. In each group, at least 20 pairs of control and knockdown cells were quantified.

(F) TMEM115 does not regulate the total and Golgi protein levels of Rab6A. Golgi membranes were isolated from the indicated cell-lines.

(G, H) TMEM115 does not regulate the Golgi localisation of Rab6A. (G) Cells stably expressing GFP-Rab6A ($WT^{GFP\text{-}Rab6A}$) were knocked down for control or TMEM115, then co-cultured. The fluorescent intensity of Golgi GFP-Rab6A was quantified in (H). Twenty-three pairs of control and knockdown cells were quantified.

(I, J) TMEM115 does not regulate Golgi Rab6 activity. (I) KO6 cells were transiently transfected with TMEM115-mNeonGreen (TMEM115-mNG), and the fluorescent intensity of Golgi of $Rab6^{GTP}$ was quantified in (J). Eighteen pairs of untransfected and transfected cells were quantified. Arrowheads in (D, G, I) indicate TMEM115 KO/KD cells; arrows in (D) indicate Rab6A KD cells.

**Figure S4 The 1-270 residues of TMEM115 are required for its Golgi localisation**

(A) The predicted structure of TMEM115 by AF3; the truncation positions are indicated.

(B) The1-270 residues of TMEM115 underpin its Golgi localisation. The 1-265 truncation of TMEM115 localises in the ER and Golgi, representing a transition of the localisation from exclusive ER (the 1-260 truncate) to exclusive Golgi (the 1-270 truncate).

**Figure S5 The cytoplasmic helix of TMEM115 is required for TMF1 Golgi localisation**

(A, B) The cytoplasmic helix of TMEM115 is essential for locating TMF1 to the Golgi. (A) KO6 cells and KO6 cells stably expressing the indicated TMEM115 constructs were co-cultured and the fluorescent intensity of Golgi TMF1 were quantified in (B). Arrowheads indicate TMEM115 KO cells.

(C, D) The cytoplasmic helix of TMEM115 is essential for preventing TMF1 from accumulating in vesicles. Total vesicles were isolated from the indicated cell lines.

**Figure S6 All the coiled-coils of TMF1 are simultaneously required for its Golgi localisation**

(A) The TMF1 deletion constructs used in this experiment. The ability of Golgi localisation of each construct is summarised to the right

(B) All the coiled coils of TMF1 are required for its Golgi localisation. Flag-tagged TMF1 truncates were stably expressed in cells.

**Figure S7 TMEM115, TMF1, and Rab6A form a ternary complex *in vitro***

(A – C) Cryo-EM, image processing workflows, and example 2D class averages of the TTR complex. Red boxed images were used as representative 2D class averages in Figure 4B.

(D) Confidence scores mapped onto the AlphaFold 3-predicted TTR complex.

**Figure S8 Assembly of the TTR complex *in vitro***

(A) Crystal structures of the human Rab6-human GCC2 complex (top) and the human Rab6A - mouse Kif20A complex (bottom).

(B, C) TMEM115 is dispensable for TMF1 and Rab6A assembly under *in vitro* controlled conditions. *In vitro* purified proteins were mixed in combination to examine their ability for forming complexes. The eluant from a mixture of the proteins was fractionated on an S6 10/300 size-exclusion column. Proteins present in fractions were analysed by SDS-PAGE and Coomassie staining

**Figure S9 Predicted structures of the TTR complex**

(A, B) Details of the TMEM115-TMF1 and TMF1-Rab6A interfaces.

(C) The residues of TMEM115, Rab6A, and TMF1 involved in TTR complex formation are conserved. Primary sequences of TMEM115, Rab6A, and TMF1 proteins across yeast, fly, mouse, and human were aligned and residues involved in the interactions are highlighted in red.

**Figure S10 Human disease mutations of TMF1 and TMEM115 affect TMF1 Golgi localisation**

(A, B) Human disease mutations of TMF1 affect its Golgi localisation and TTR complex formation. Flag-TMF1 mutants were transiently expressed in RPE1 cells, followed by microscopy (A) or IP assays (B).

(C, D) Human disease mutations of TMEM115 affect TMF1 Golgi localisation and TTR complex formation. TMEM115 KO and the KO cells stably expressing the indicated TMEM115 human disease mutants were co-cultured, followed by microscopy (C) and IP assays (D).

(E) Principle of the mitochondria anchoring-away experiment.

**Figure S11 Rabs on their own are inefficient in locating effector proteins**

(A, B) Mitochondrial Rab6A$^{GTP}$ on its own is inefficient in relocating TMF1and BICD2. Cells were transiently over-expressed with Rab6A$^{Q72L}$-HA-MaoB, left untreated (UNT) or treated with BFA.

(C) Mitochondrial Rab5A$^{GTP}$ on its own is inefficient in relocating EEA1. Cells were transiently over-expressed with Rab5A$^{Q79L}$-HA-MaoB, left untreated or treated with nocodazole (Noc). Arrowheads indicate cells expressing extremely high levels of the mitochondrial Rabs could occasionally relocate effector proteins.

**Figure S12 The TMEM115-TMF1 axis regulates Golgi proteome**

(A) Proteomics correlation analysis across two TMF1 KD datasets. Only consistently altered protein hits were plotted.

(B) Proteomics correlation analysis across two TMEM115 KD and two TMF1 KD datasets. Only consistently altered protein hits were plotted.

**Figure S13 Generation and phenotyping of the *TMEM115* and *CG9536* mutant animals**

(A) Scheme of generating the *TMEM115* KO mice. Details are described in *Materials and Methods.*

(B, C) TMEM115 KO mice show less weight after birth and struggle to gain weight over time. (B) Newborn mice were genotyped and weighed. (C) Weight gain of the different genotypes during the first 25 days of life. No KO animals lived more than 22 days. Data is shown as mean ± SDs.

(D) Scheme of generating the *CG9536* KO flies. Details are described in *Materials and Methods.*

### Figure S1

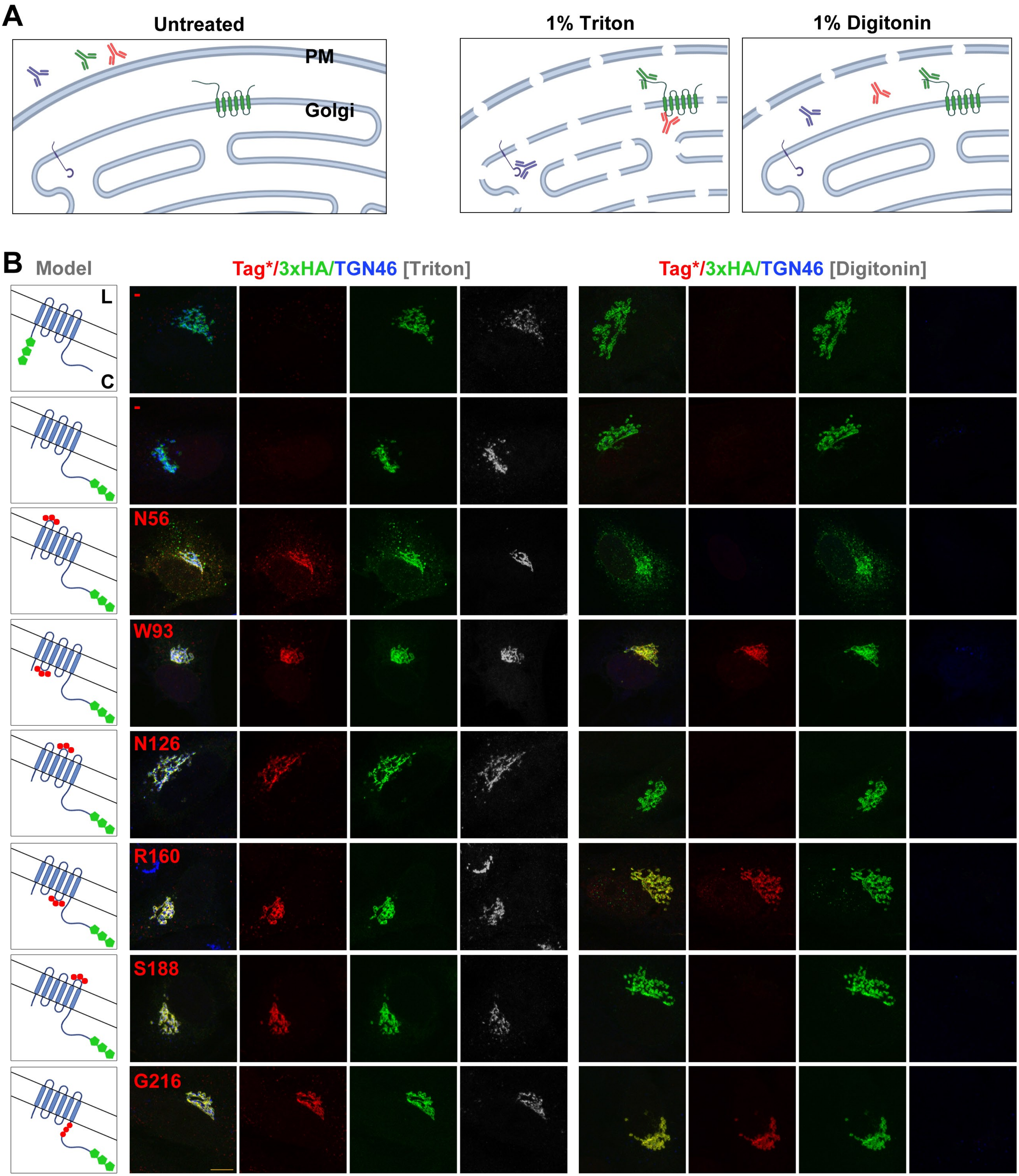
A
Untreated
PM
Golgi
1% Triton
1% Digitonin
B
Model
L
C
Tag*/3xHA/TGN46 [Triton]
Tag*/3xHA/TGN46 [Digitonin]
-
-
N56
W93
N126
R160
S188
G216

### Figure S2

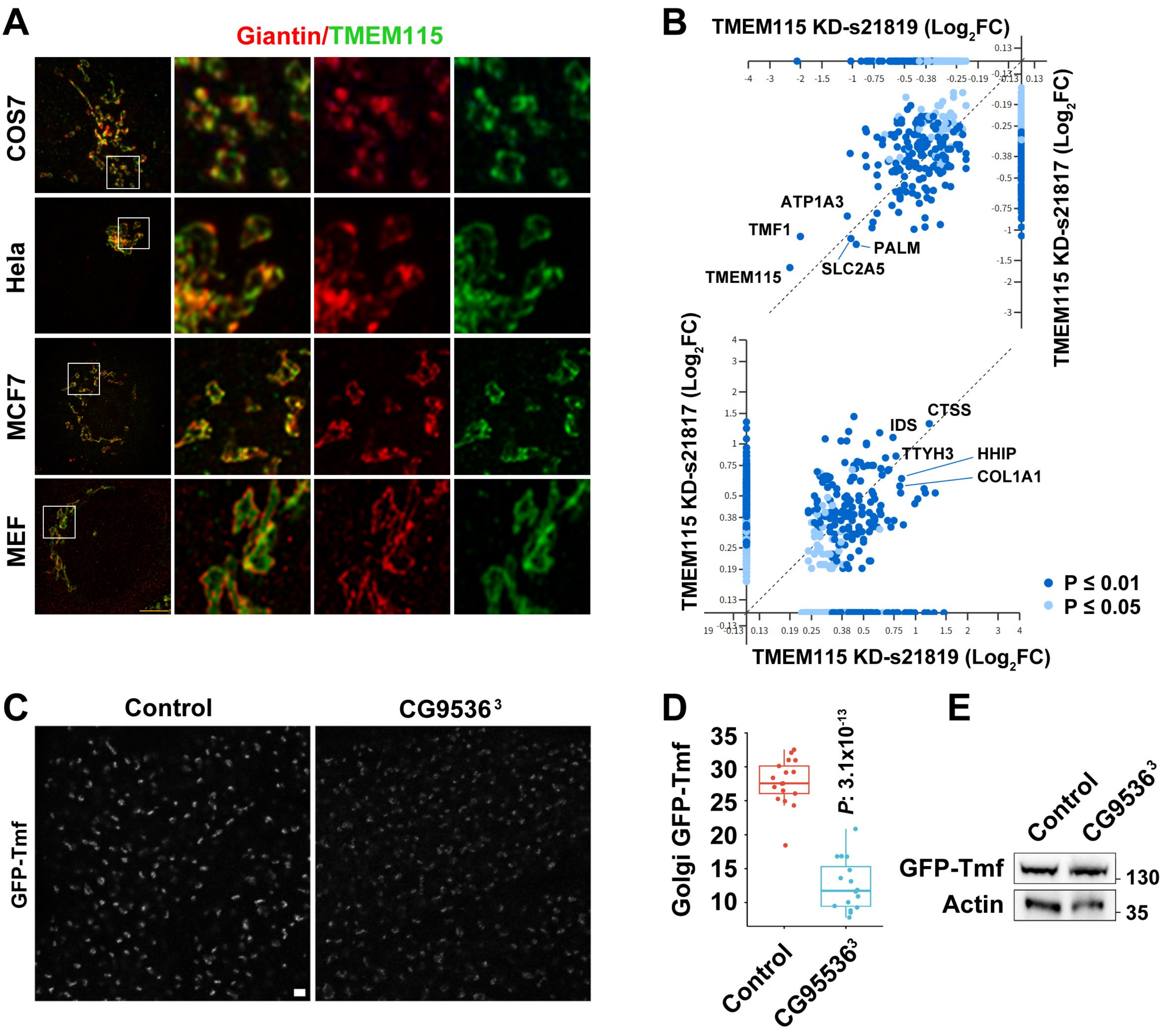
A
Giantin/TMEM115
COS7
Hela
MCF7
MEF
B
TMEM115 KD-s21819 (Log2FC)
TMEM115 KD-s21817 (Log2FC)
ATP1A3
TMF1
PALM
SLC2A5
TMEM115
CTSS
IDS
TTYH3
HHIP
COL1A1
P ≤ 0.01
P ≤ 0.05
C
Control
CG9536 3
GFP-Tmf
D
Golgi GFP-Tmf
P: 3.1x10-13
Control
CG9536 3
E
Control
CG9536 3
GFP-Tmf
Actin
130
35

### Figure S3

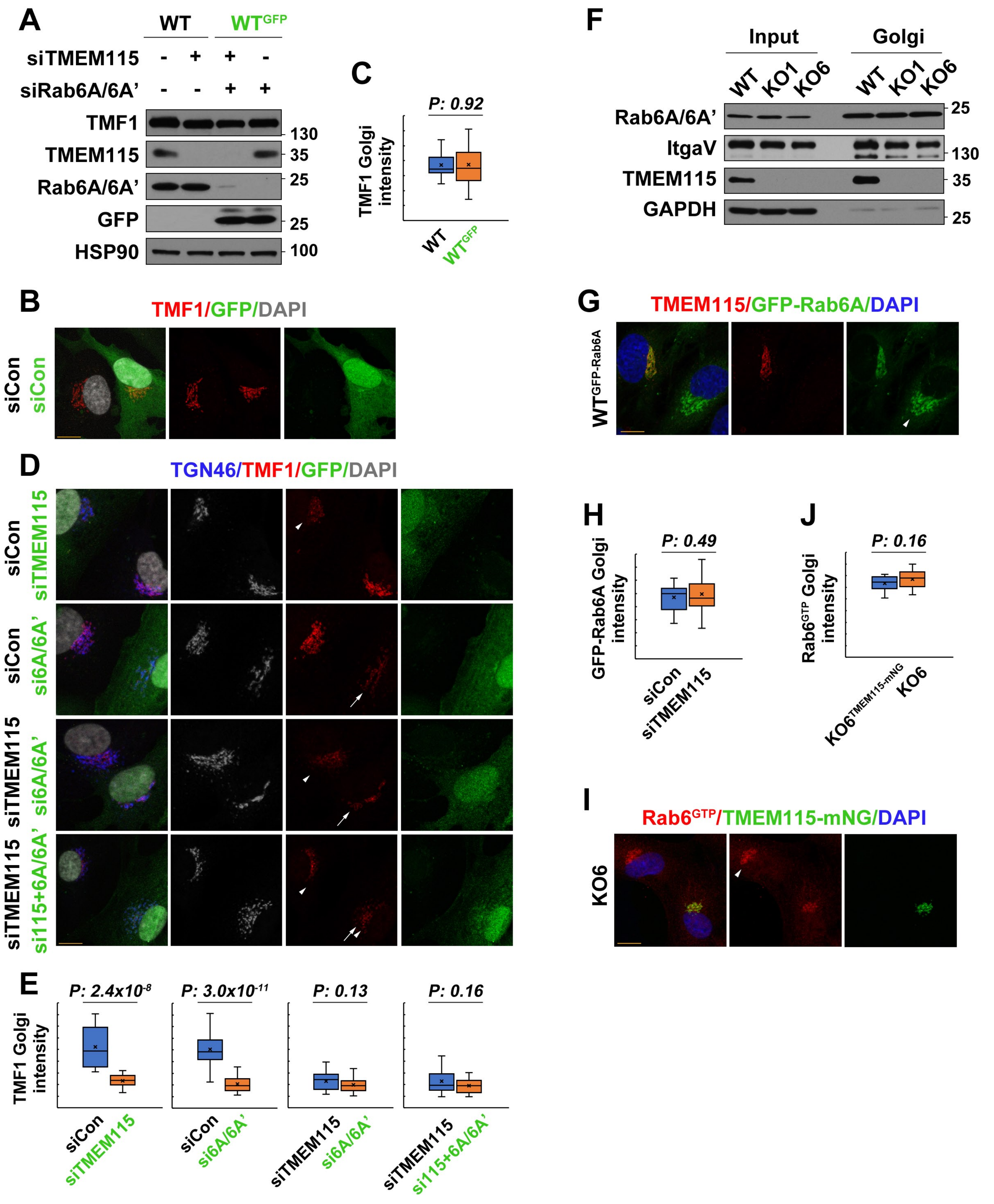

A
WT
WTGFP
siTMEM115 - + + -
siRab6A/6A' - - + +
TMF1
TMEM115
Rab6A/6A'
GFP
HSP90
130
35
25
25
100
C
P: 0.92
TMF1 Golgi intensity
WT
WTGFP
F
Input
Golgi
WT KO1 KO6 WT KO1 KO6
Rab6A/6A'
ItgaV
TMEM115
GAPDH
25
130
35
25
B
TMF1/GFP/DAPI
siCon siCon
G
TMEM115/GFP-Rab6A/DAPI
WTGFP-Rab6A
D
TGN46/TMF1/GFP/DAPI
siCon siTMEM115
siCon si6A/6A'
siTMEM115 si6A/6A'
siTMEM115 si115+6A/6A'
H
P: 0.49
GFP-Rab6A Golgi intensity
siCon
siTMEM115
J
P: 0.16
Rab6GTP Golgi intensity
KO6TMEM115-mNG
KO6
I
Rab6GTP/TMEM115-mNG/DAPI
KO6
E
TMF1 Golgi intensity
P: 2.4x10-8
P: 3.0x10-11
P: 0.13
P: 0.16
siCon siTMEM115
siCon si6A/6A'
siTMEM115 si6A/6A'
siTMEM115 si115+6A/6A'

### Figure S4

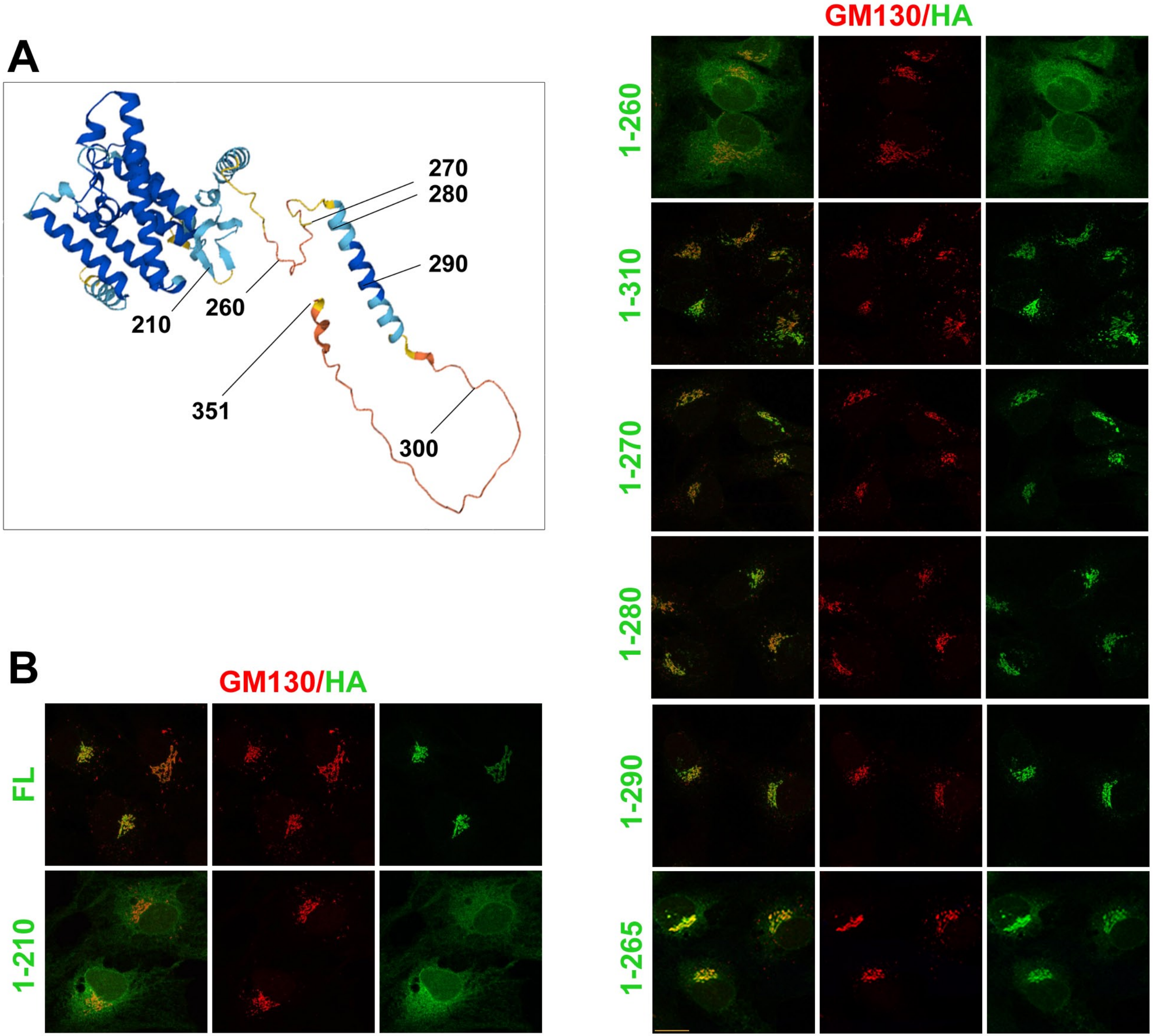
A
270
280
290
260
210
351
300
B
GM130/HA
FL
1-210
GM130/HA
1-260
1-310
1-270
1-280
1-290
1-265

### Figure S5

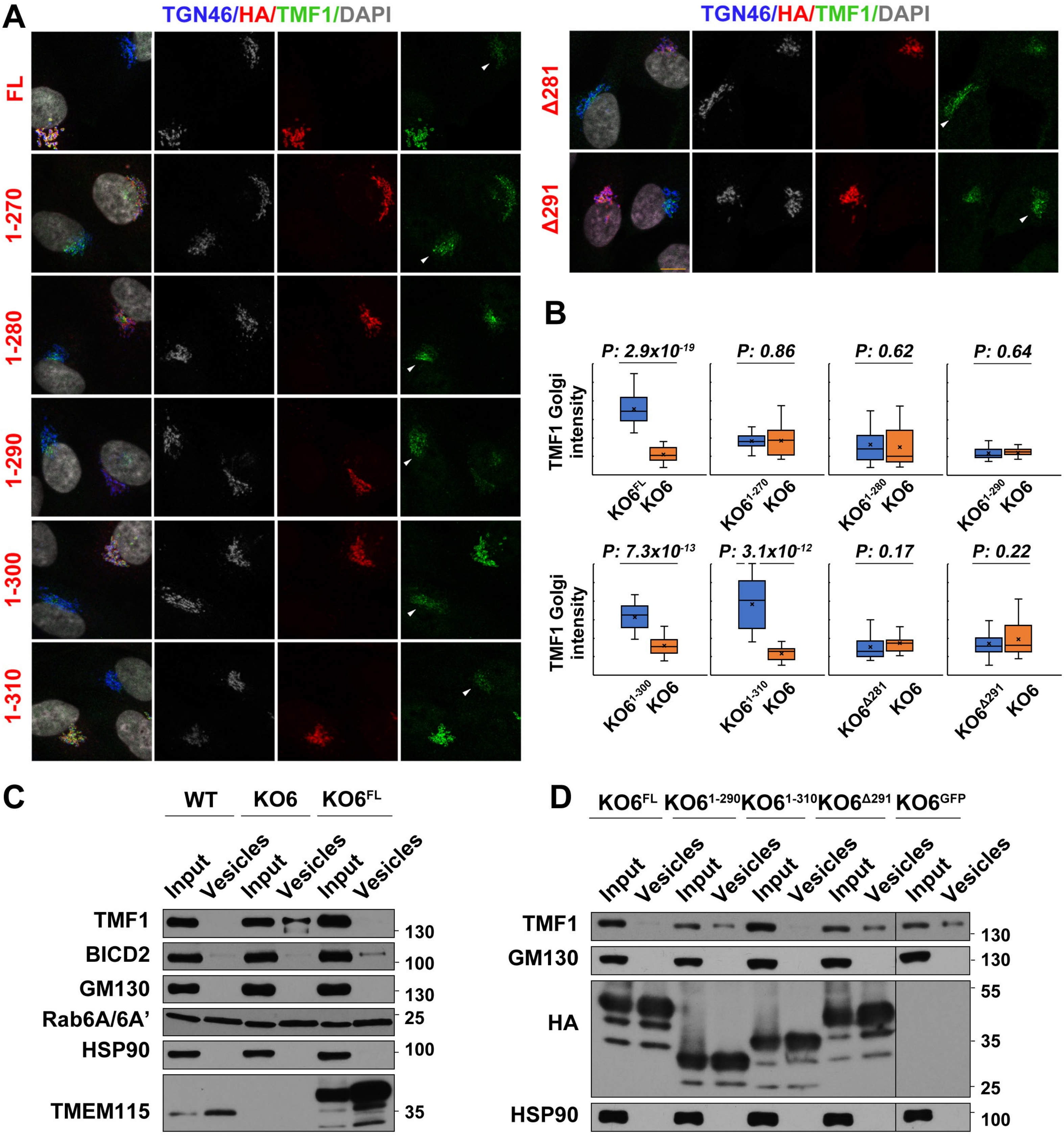

A
TGN46/HA/TMF1/DAPI
FL
1-270
1-280
1-290
1-300
1-310
Δ281
Δ291
B
TMF1 Golgi intensity
P: 2.9x10-19
P: 0.86
P: 0.62
P: 0.64
KO6FL KO6
KO61-270 KO6
KO61-280 KO6
KO61-290 KO6
P: 7.3x10-13
P: 3.1x10-12
P: 0.17
P: 0.22
KO61-300 KO6
KO61-310 KO6
KO6Δ281 KO6
KO6Δ291 KO6
C
WT
KO6
KO6FL
Input
Vesicles
TMF1
BICD2
GM130
Rab6A/6A'
HSP90
TMEM115
130
100
130
25
100
35
D
KO6FL
KO61-290
KO61-310
KO6Δ291
KO6GFP
Input
Vesicles
TMF1
GM130
HA
HSP90
130
130
55
35
25
100

### Figure S6

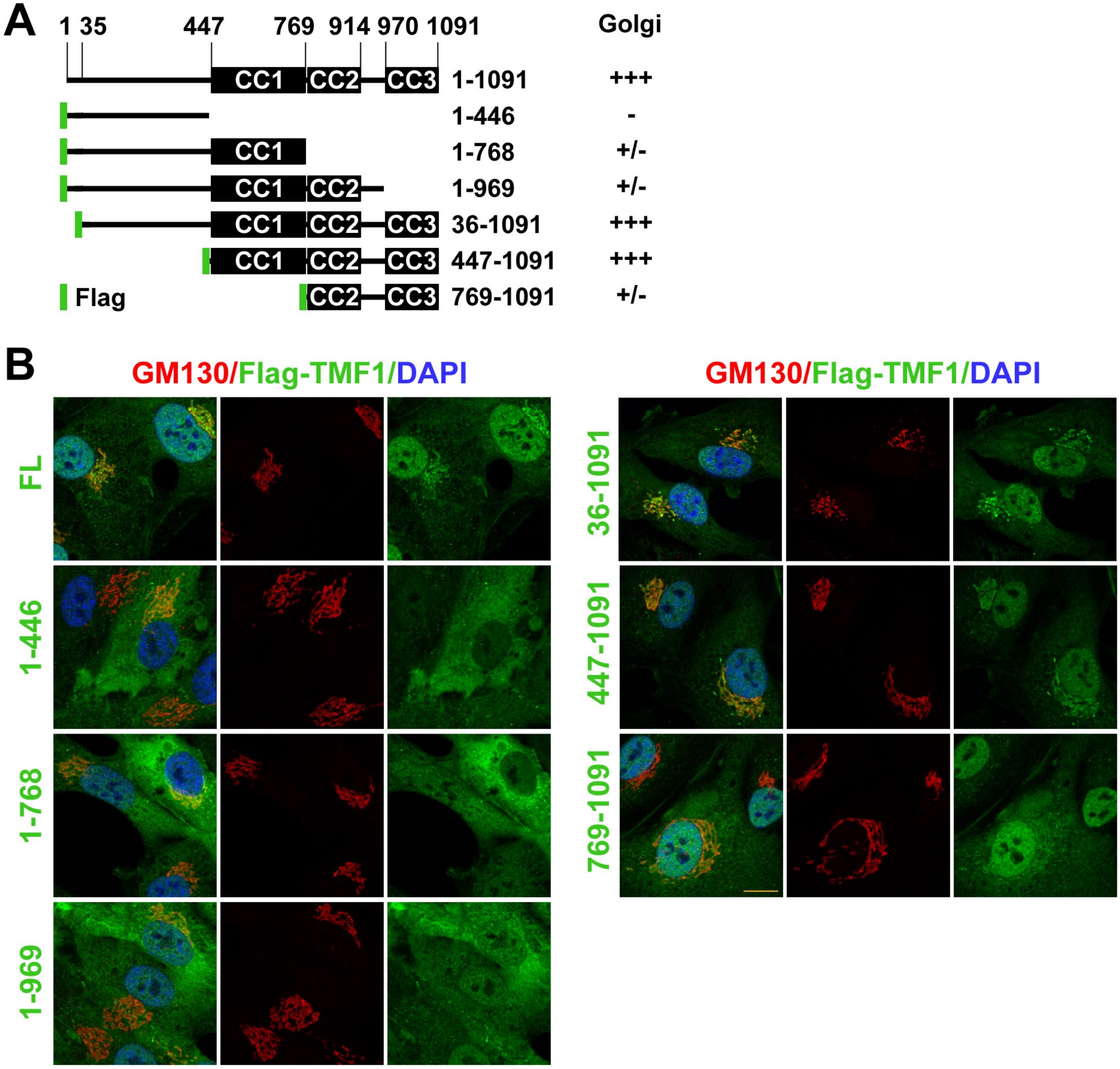
A
1 35 447 769 914 970 1091
Golgi
CC1 CC2 CC3 1-1091 +++
1-446 -
CC1 1-768 +/-
CC1 CC2 1-969 +/-
CC1 CC2 CC3 36-1091 +++
CC1 CC2 CC3 447-1091 +++
Flag CC2 CC3 769-1091 +/-
B
GM130/Flag-TMF1/DAPI
FL
1-446
1-768
1-969
GM130/Flag-TMF1/DAPI
36-1091
447-1091
769-1091

### Figure S7

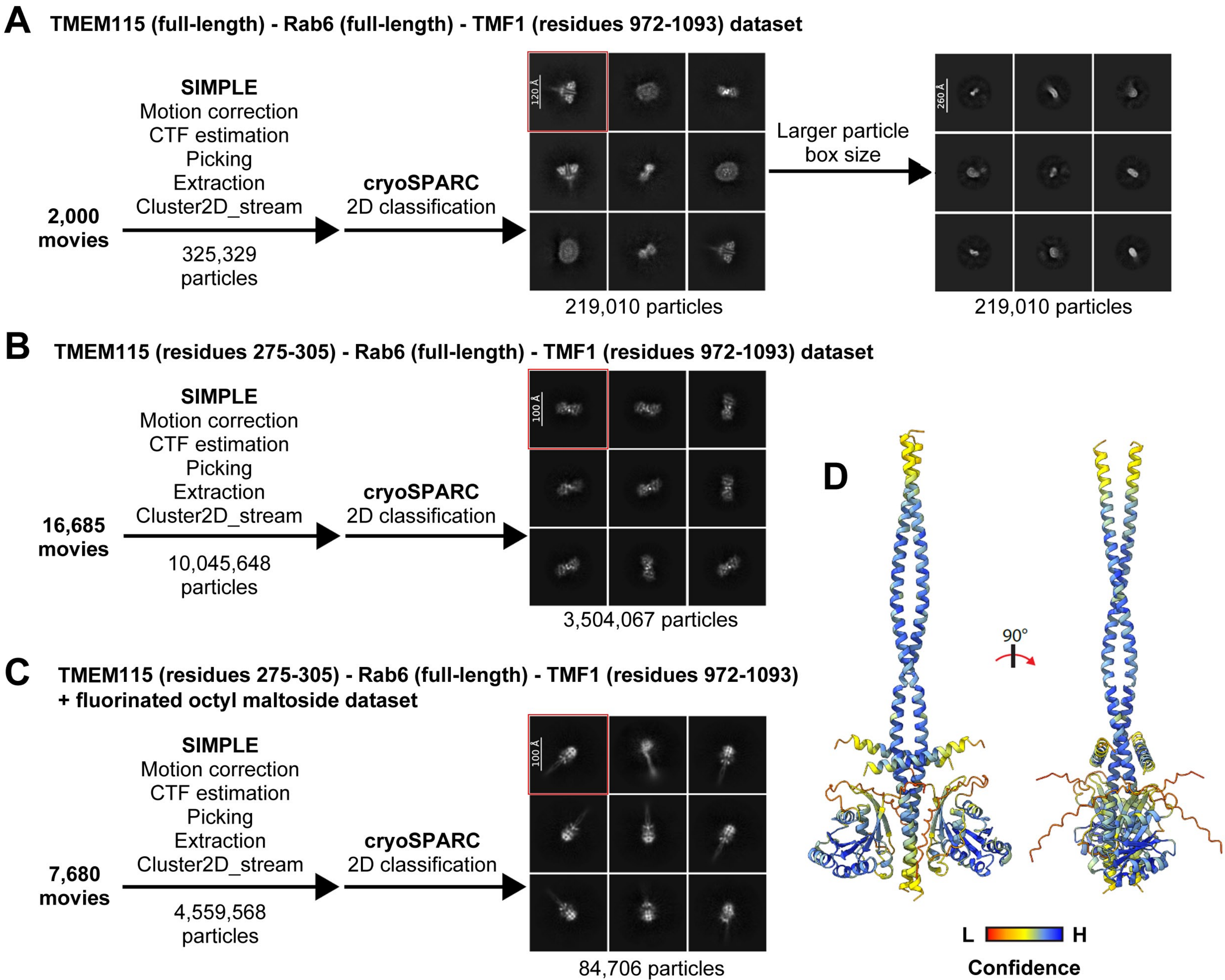
A
TMEM115 (full-length) - Rab6 (full-length) - TMF1 (residues 972-1093) dataset
2,000 movies
SIMPLE
Motion correction
CTF estimation
Picking
Extraction
Cluster2D_stream
325,329 particles
cryoSPARC
2D classification
120 Å
219,010 particles
Larger particle box size
260 Å
219,010 particles
B
TMEM115 (residues 275-305) - Rab6 (full-length) - TMF1 (residues 972-1093) dataset
16,685 movies
SIMPLE
Motion correction
CTF estimation
Picking
Extraction
Cluster2D_stream
10,045,648 particles
cryoSPARC
2D classification
100 Å
3,504,067 particles
C
TMEM115 (residues 275-305) - Rab6 (full-length) - TMF1 (residues 972-1093) + fluorinated octyl maltoside dataset
7,680 movies
SIMPLE
Motion correction
CTF estimation
Picking
Extraction
Cluster2D_stream
4,559,568 particles
cryoSPARC
2D classification
100 Å
84,706 particles
D
90°
L
H
Confidence

#### Figure S8

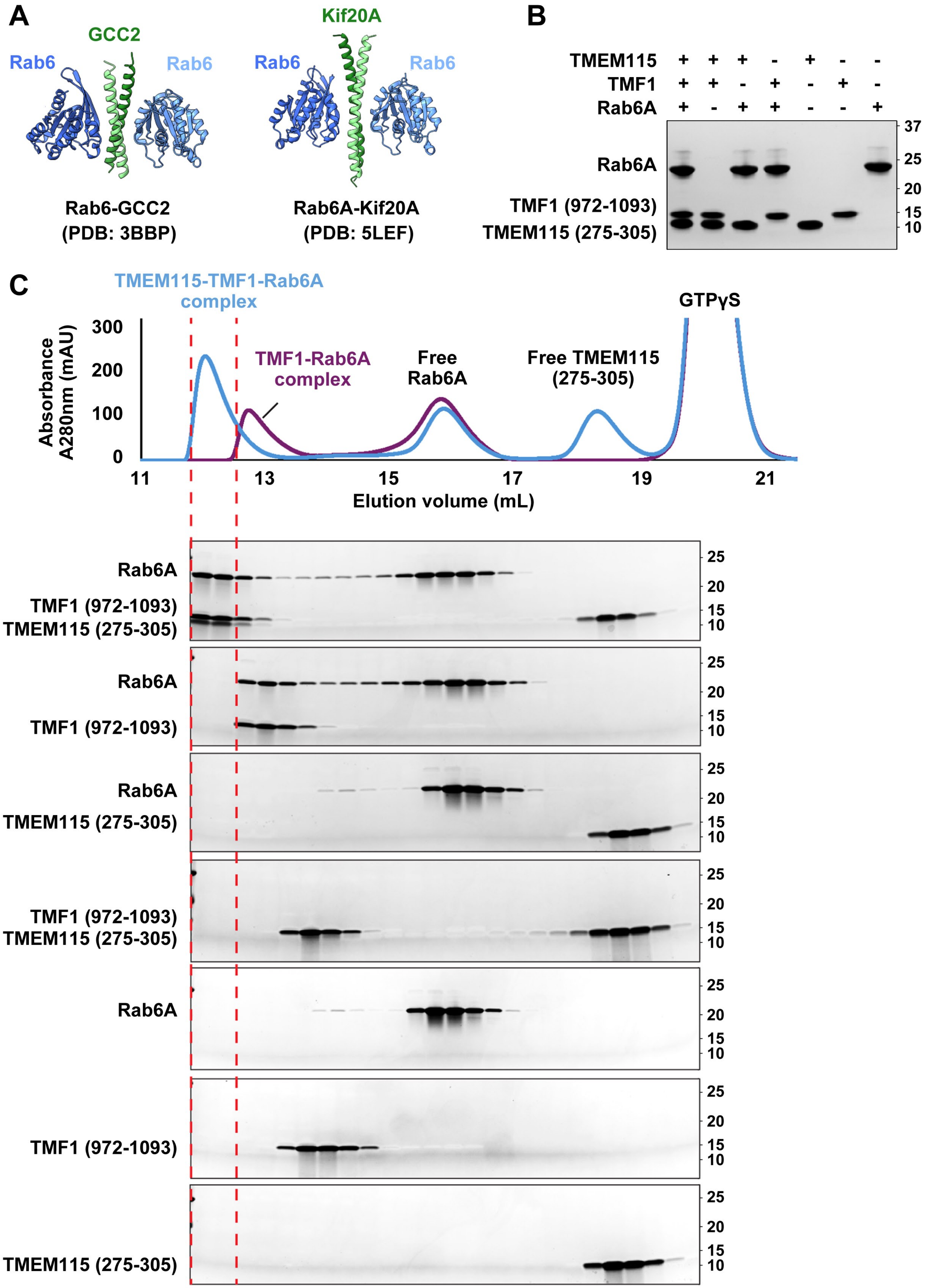

A
GCC2
Rab6
Rab6
Rab6-GCC2
(PDB: 3BBP)
Kif20A
Rab6
Rab6
Rab6A-Kif20A
(PDB: 5LEF)
B
TMEM115 + + + - + - -
TMF1 + + - + - + -
Rab6A + - + + - - +
Rab6A
TMF1 (972-1093)
TMEM115 (275-305)
37
25
20
15
10
C
TMEM115-TMF1-Rab6A complex
TMF1-Rab6A complex
Free Rab6A
Free TMEM115 (275-305)
GTPγS
Absorbance A280nm (mAU)
300
200
100
0
11
13
15
17
19
21
Elution volume (mL)
Rab6A
TMF1 (972-1093)
TMEM115 (275-305)
25
20
15
10

### Figure S9

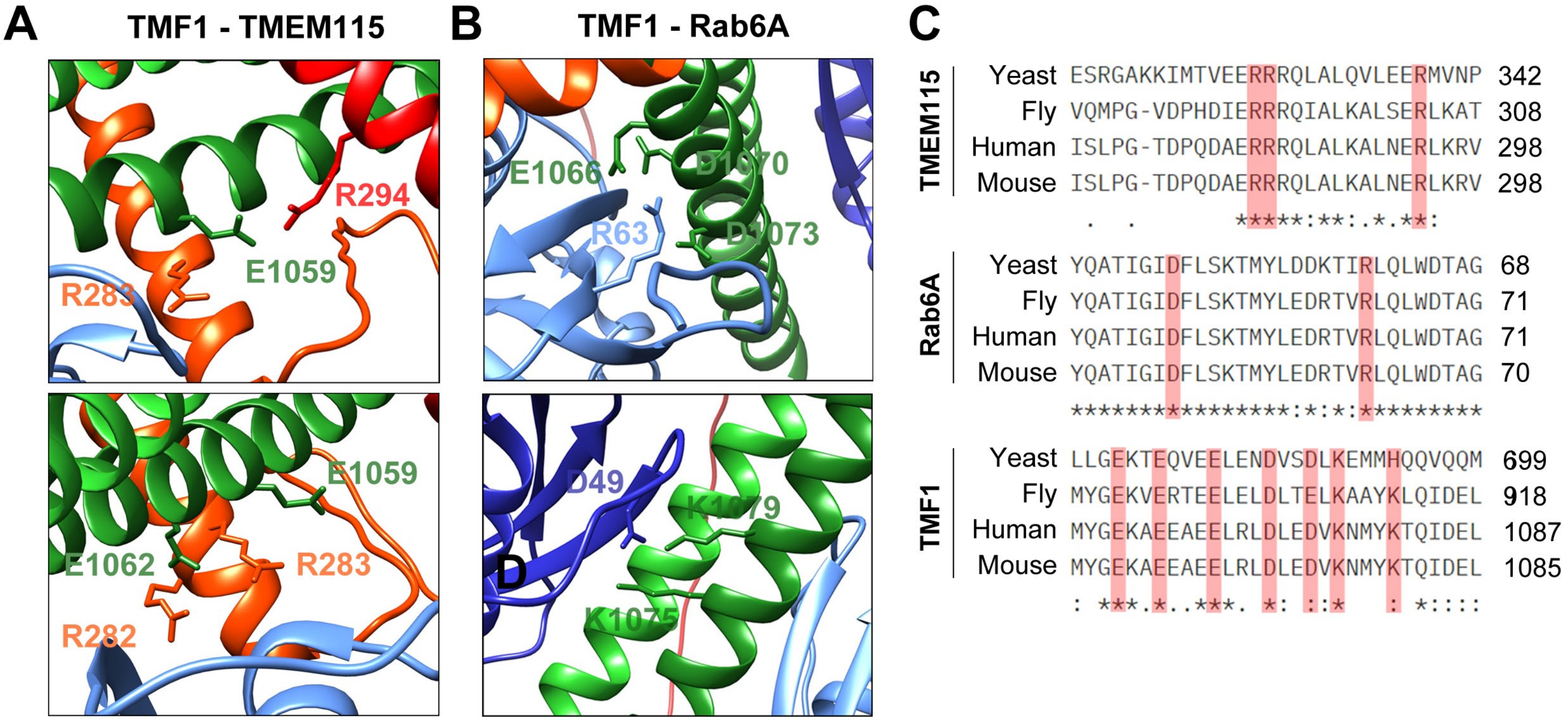

A
TMF1 - TMEM115
R294
E1059
R283
E1059
E1062
R283
R282
B
TMF1 - Rab6A
E1066
D1070
R63
D1073
D49
K1079
K1075
D
C
TMEM115
Yeast ESRGAKKIMTVEERRRQLALQVLEERMVNP 342
Fly VQMPG-VDPHDIERRRQIALKALSERLKAT 308
Human ISLPG-TDPQDAERRRQLALKALNERLKRV 298
Mouse ISLPG-TDPQDAERRRQLALKALNERLKRV 298
Rab6A
Yeast YQATIGIDFLSKTMYLDDKTIRLQLWDTAG 68
Fly YQATIGIDFLSKTMYLEDRTVRLQLWDTAG 71
Human YQATIGIDFLSKTMYLEDRTVRLQLWDTAG 71
Mouse YQATIGIDFLSKTMYLEDRTVRLQLWDTAG 70
TMF1
Yeast LLGEKTEQVEELENDVSDLKEMMHQQVQQM 699
Fly MYGEKVERTEELELDLTELKAAYKLQIDEL 918
Human MYGEKAEEAEELRLDLEDVKNMYKTQIDEL 1087
Mouse MYGEKAEEAEELRLDLEDVKNMYKTQIDEL 1085

### Figure S10

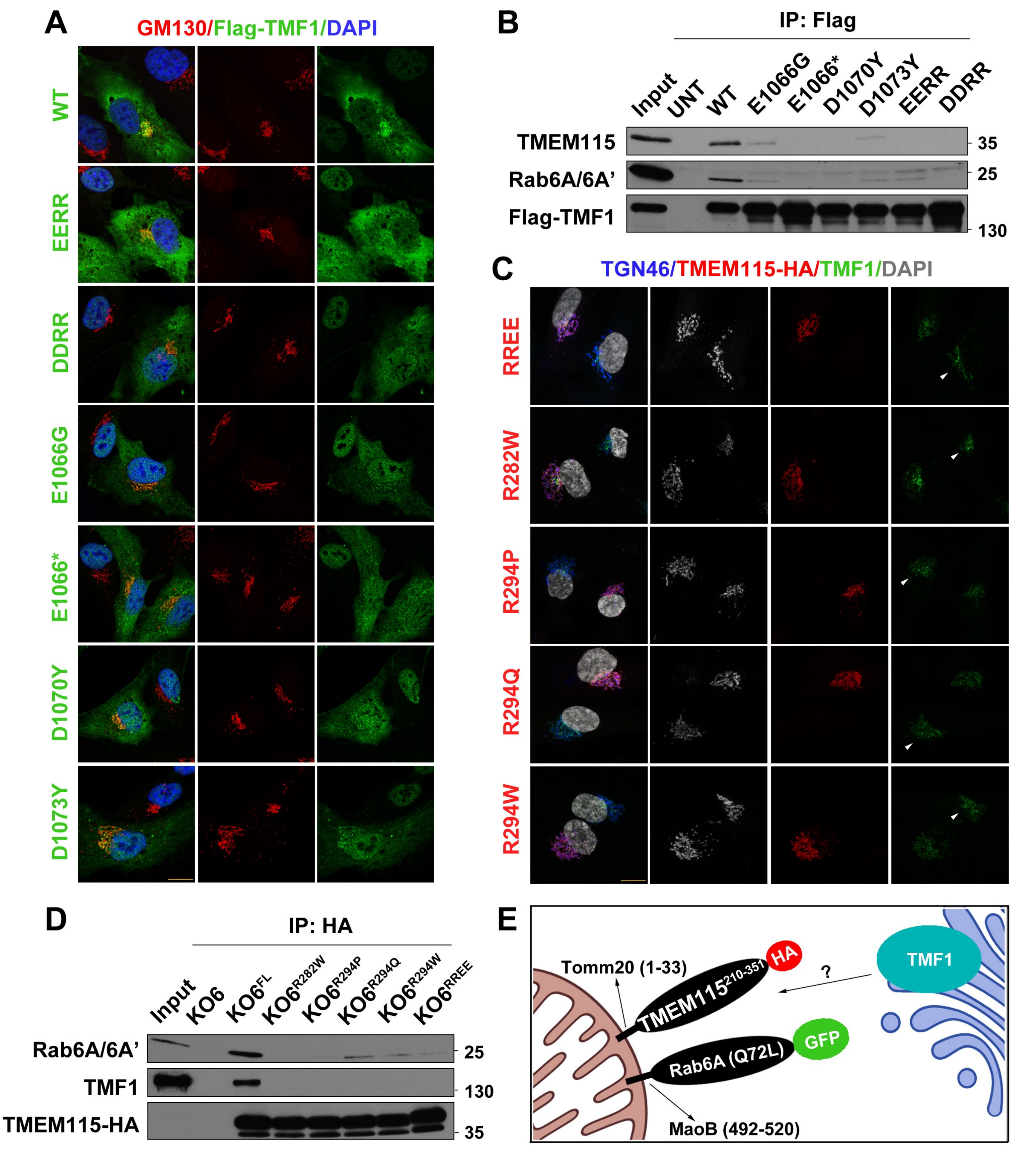
A
GM130/Flag-TMF1/DAPI
WT
EERR
DDRR
E1066G
E1066*
D1070Y
D1073Y
B
IP: Flag
Input
UNT
WT
E1066G
E1066*
D1070Y
D1073Y
EERR
DDRR
TMEM115
Rab6A/6A'
Flag-TMF1
35
25
130
C
TGN46/TMEM115-HA/TMF1/DAPI
RREE
R282W
R294P
R294Q
R294W
D
IP: HA
Input
KO6
KO6FL
KO6R282W
KO6R294P
KO6R294Q
KO6R294W
KO6RREE
Rab6A/6A'
TMF1
TMEM115-HA
25
130
35
E
Tomm20 (1-33)
TMEM115210-351
HA
?
TMF1
Rab6A (Q72L)
GFP
MaoB (492-520)

### Figure S11

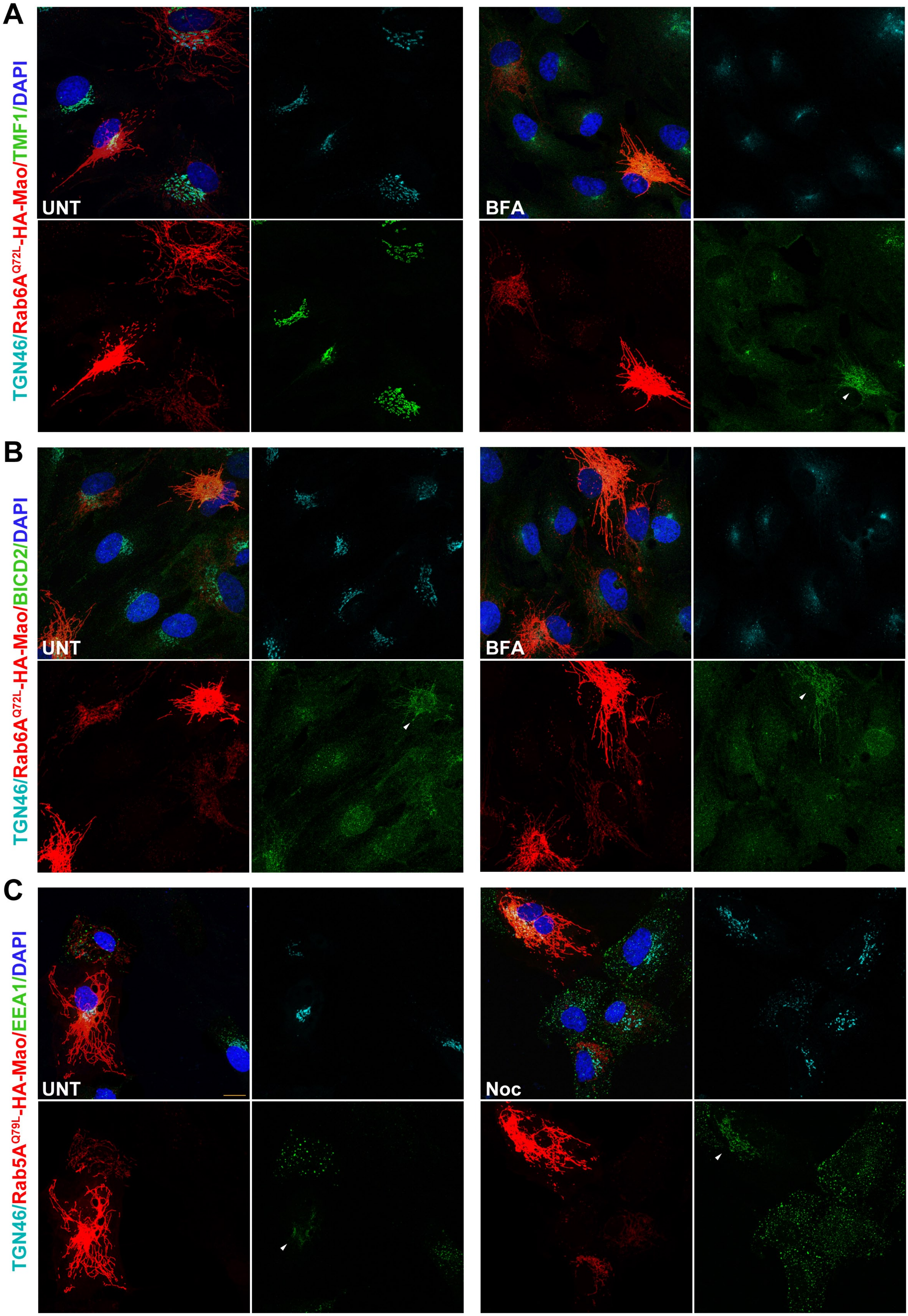
A
TGN46/Rab6A$^{Q72L}$-HA-Mao/TMF1/DAPI
UNT
BFA
B
TGN46/Rab6A$^{Q72L}$-HA-Mao/BICD2/DAPI
UNT
BFA
C
TGN46/Rab5A$^{Q79L}$-HA-Mao/EEA1/DAPI
UNT
Noc

### Figure S12

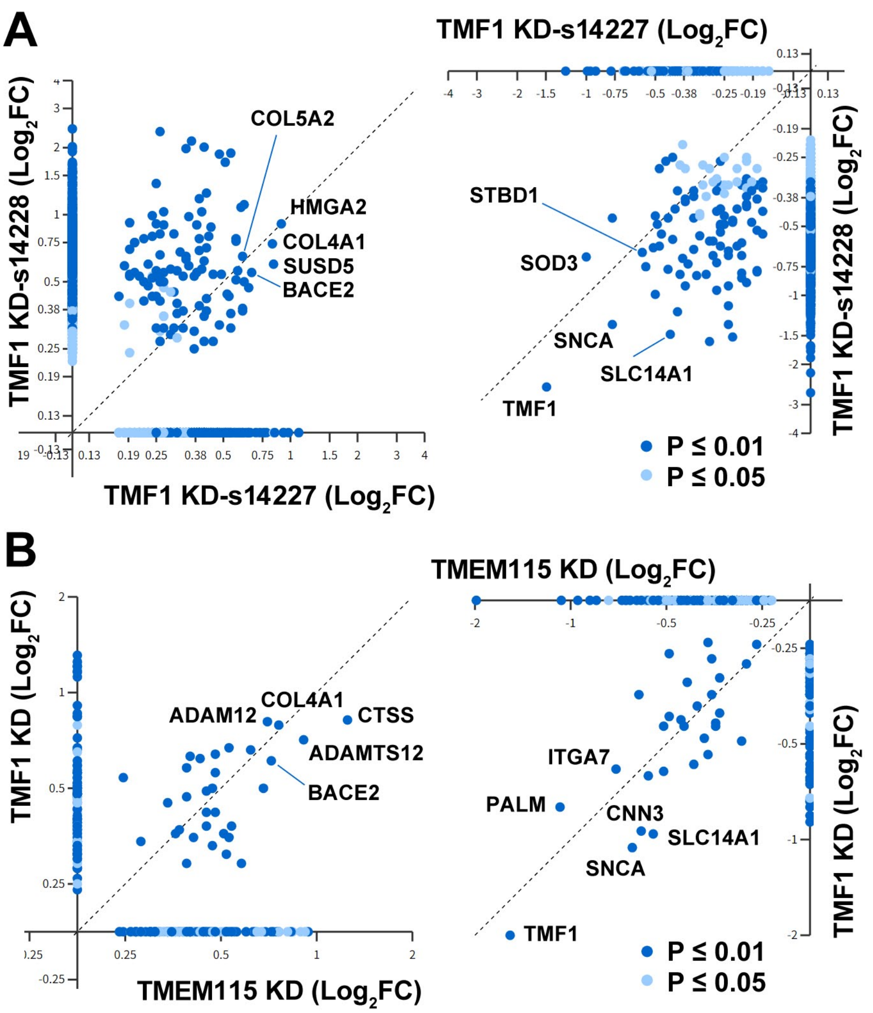
A
TMF1 KD-s14227 (Log2FC)
TMF1 KD-s14228 (Log2FC)
COL5A2
HMGA2
COL4A1
SUSD5
BACE2
STBD1
SOD3
SNCA
SLC14A1
TMF1
P ≤ 0.01
P ≤ 0.05
B
TMEM115 KD (Log2FC)
TMF1 KD (Log2FC)
ADAM12
COL4A1
CTSS
ADAMTS12
BACE2
ITGA7
PALM
CNN3
SLC14A1
SNCA
TMF1
P ≤ 0.01
P ≤ 0.05

### Figure S13

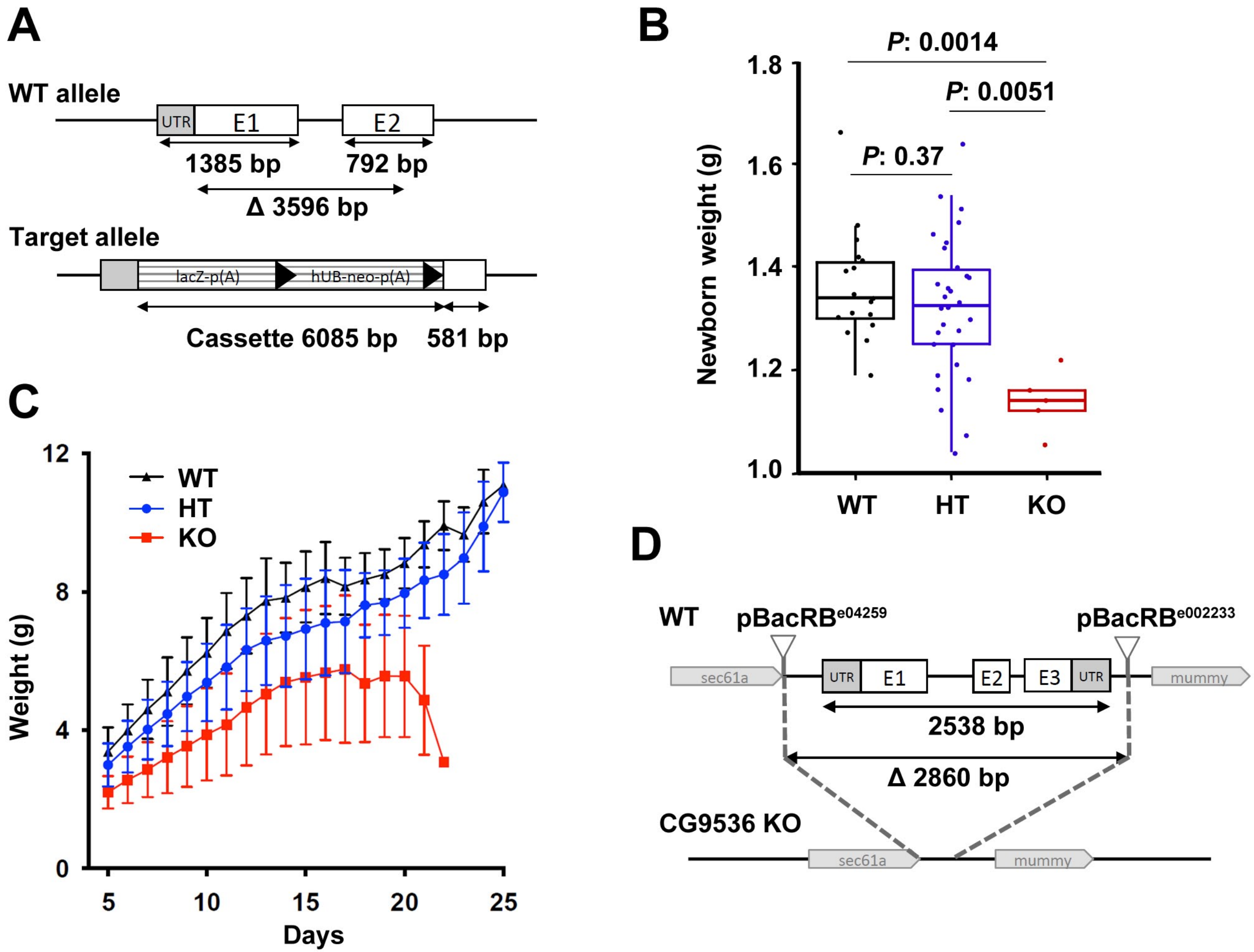

A
WT allele
UTR
E1
E2
1385 bp
792 bp
Δ 3596 bp
Target allele
lacZ-p(A)
hUB-neo-p(A)
Cassette 6085 bp
581 bp
B
P: 0.0014
P: 0.0051
P: 0.37
Newborn weight (g)
1.8
1.6
1.4
1.2
1.0
WT
HT
KO
C
WT
HT
KO
Weight (g)
12
8
4
0
5
10
15
20
25
Days
D
WT
pBacRB^e04259
pBacRB^e002233
sec61a
UTR
E1
E2
E3
UTR
mummy
2538 bp
Δ 2860 bp
CG9536 KO
sec61a
mummy
